# Par3/Bazooka binds NICD and promotes Notch signalling during *Drosophila* development

**DOI:** 10.1101/2022.05.24.493322

**Authors:** Jun Wu, Neeta Bala Tannan, Linh T. Vuong, Yildiz Koca, Giovanna M. Collu, Marek Mlodzik

## Abstract

The conserved *bazooka (baz/par3*) gene acts as a key regulator of asymmetrical cell divisions across the animal kingdom. Associated Par3/Baz-Par6-aPKC protein complexes are also well known for their role in the establishment of apical/basal cell polarity in epithelial cells. Here we define a novel, positive function of Baz/Par3 in the Notch pathway. Using *Drosophila* wing and eye development, we demonstrate that Baz is required for Notch signaling activity and optimal transcriptional activation of Notch target genes. Baz appears to act independently of aPKC in these contexts, as knockdown of *aPKC* does not cause *Notch* loss-of-function phenotypes. Using transgenic Notch constructs, our data positions Baz activity downstream of activating Notch cleavage steps and upstream of Su(H)/CSL transcription factor complex activity on Notch target genes. We demonstrate a biochemical interaction between NICD and Baz, suggesting that Baz is required for NICD activity before NICD binds to Su(H). Taken together, our data define a novel role of the polarity protein Baz/Par3, as a positive and direct regulator of Notch signaling through its interaction with NICD.

## Introduction

Cell polarity is a critical feature of most cell types, particularly epithelial cell sheets, which display obligate apical-basal (A/B) polarity that is required for their barrier function and vectorial secretion. Many proteins and mechanisms that govern the establishment and maintenance of cell polarity are evolutionarily conserved and were discovered genetically in *C. elegans* and *Drosophila* (reviewed in (Goldstein and Macara, 2007; St Johnston and Sanson, 2011; Tepass, 2012)). Similarly, stem cells often display polarized features using the same factors, which are critical for stem cell maintenance and for progenitor cell fates, with cell contents being asymmetrically inherited by the two daughter cells (e.g. rev. in (Neumuller and Knoblich, 2009; Schweisguth, 2015)). The protein complexes that mediate polarization are also shared among these distinct instances of cell polarization.

The conserved *par3* gene, known as *bazooka/baz* in *Drosophila*, was originally identified in *C. elegans*, where it acts as a key regulator of asymmetrical cell divisions in the early embryo (Etemad-Moghadam et al., 1995). It generally acts together with Par6 and aPKC as a regulatory complex of many asymmetric cell divisions across the animal kingdom (reviewed in (Neumuller and Knoblich, 2009; Schweisguth, 2015)). This Par3/Baz-Par6-aPKC complex is also well known for its conserved role in the establishment of A/B polarity in epithelial cells (Suzuki and Ohno, 2006). Our interest in the A/B epithelial cell polarity proteins originates from the fact that several of these have also been linked to planar cell polarity (PCP), that is polarity within the epithelial sheet, orthogonal to the A/B axis (Humphries and Mlodzik, 2018). For example aPKC can associate with Frizzled, and Scribble (another A/B polarity protein) associates with the PCP protein Van Gogh/Vang (Vangl in vertebrates) (Courbard et al., 2009; Djiane et al., 2005). aPKC and Scribble thus contribute to the stability and asymmetric segregation of the core PCP complexes and establishment of planar polarity within epithelial cell sheets. To systematically examine whether additional A/B epithelial polarity factors can play a role in PCP establishment, we tested members of these protein families for PCP associated knockdown phenotypes during *Drosophila* wing development. Surprisingly when we examined phenotypes of *baz/par3* knockdown in this context, we observed a *Notch* mutant-like phenotype of loss of wing margin tissue. Here we focus on investigating a specific role for Baz in regulating Notch signaling.

*Notch (N)* was originally identified in *Drosophila* based on its haplo-insufficient, dominant wing-notching phenotype of loss of tissue at the dorso-ventral border of the wing (Guruharsha et al., 2012; Mohr, 1919; Southgate et al., 2015). *Notch* encodes a single-pass transmembrane receptor protein containing EGF-like repeats in its extracellular domain (Wharton et al., 1985) and a conserved intracellular domain (NICD) that, upon cleavage, can translocate to the nucleus and function as transcriptional co-activator (reviewed in (Andersson et al., 2011; Bray, 2016; Henrique and Schweisguth, 2019; Kopan and Ilagan, 2009; Wilson and Radtke, 2006)). Many studies demonstrated that the highly conserved Notch receptor family plays important roles during the development and homeostasis of most tissues through cell-cell communication and cellular signaling, controlling many target genes that can be either common or context dependent (reviewed in (Andersson et al., 2011; Bray, 2016; Henrique and Schweisguth, 2019; Kopan and Ilagan, 2009; Wilson and Radtke, 2006) (Siebel and Lendahl, 2017)). Misregulation of N-signaling is associated with many human diseases, including cancer (Siebel and Lendahl, 2017). The Notch receptor is activated by the binding of ligands of the Delta (Dl) or Serrate/Jagged (Ser) families from neighboring cells to trigger juxtamebrane cleavage events to release NICD (Andersson et al., 2011; Bray, 2016; Struhl and Greenwald, 1999). NICD can then translocate into the nucleus, where it interacts with transcription factors of the Suppressor of Hairless (Su(H)/CSL) family, and co-activators, including Mastermind, to activate target gene expression (Bray, 2016). Notch signaling is subject to many regulatory steps that have been and continue to be identified and characterized in *Drosophila*, ranging from receptor and ligand trafficking, receptor glycosylation, membrane lipid composition, receptor cleavage, and protein stability regulators (Bala Tannan et al., 2018; Kandachar and Roegiers, 2012; Kidd et al., 1998; Medina-Yáñez et al., 2020; Rana and Haltiwanger, 2011; Shilo and Schejter, 2011; Singh and Mlodzik, 2012; Struhl and Greenwald, 1999; Weber et al., 2003) to specific nuclear events at the transcriptional end of pathway activation (rev. in (Andersson et al., 2011; Bray, 2016; Henrique and Schweisguth, 2019; Kopan and Ilagan, 2009; Wilson and Radtke, 2006)).

Previous work proposed an indirect role for Baz/Par3 in regulating Notch signaling during asymmetric cell divisions in *Drosophila*. During asymmetric cell division of the sensory organ precursors (SOPs), specific protein complexes segregate to opposite sides of the cell cortex in each dividing SOP cell, on one side the Baz/Par3-Par6-aPKC complex and on the other the Gai-Pins-Mud-Dlg-Numb-Neur complex. The two daughter cells that result from this cell division adopt distinct cell fates due to the asymmetric distribution of the above complexes: with one daughter cell inheriting Baz/Par3-Par6-aPKC, and the other the Gai-Pins-Mud-Dlg-Numb-Neur complex (Schweisguth, 2015). Notably, the asymmetric segregation of these complexes causes an asymmetric activation of Notch signaling, as Numb is known to repress N-signaling by reducing Notch levels at the cell surface, either through internalization or inhibition of recycling (Schweisguth, 2015). In a similar scenario, in the context of intestinal stem cell (ISC) number regulation, Baz is also required within the asymmetric apical-basal polarity complex segregation to regulate Notch activity (Wu et al., 2023). As such, an indirect role of Baz/Par3 in the regulation of N-signaling has been proposed, acting through the promotion of the asymmetric segregation of the respective apical-basal complexes to allow Notch activity in the Baz-containing daughter cells. However, due to this indirect role of Baz/Par3 in the pathway, such model systems are not suited to address potential direct roles of Baz/Par3 in Notch-signaling.

Using *Drosophila* wing and eye development, we define a novel, direct and positive function of Baz/Par3 in Notch signaling. We demonstrate that Baz is required for Notch signaling activity and activation of Notch transcriptional target genes. Baz appears to act independently of aPKC in these contexts, as knockdown of *aPKC* causes PCP defects and not Notch-associated phenotypes. Our data indicate that Baz acts the level of NICD, the cleaved active Notch fragment, but upstream of the Su(H)-NICD-Mam activation complex. We further demonstrate a direct biochemical interaction between NICD and Baz. Taken together, our data define a novel role of the polarity protein Baz/Par3, as a positive regulator of Notch signaling through its interaction with NICD.

## Results

### Reduction of *baz/par3* causes defects similar to *Notch* loss-of-function phenotypes

To initially assess potential functions of Baz/Par3-Par6-aPKC factors in PCP we expressed two independent non-overlapping RNAi (IR) lines for each gene in the posterior compartment of developing wing discs using the *en-Gal4* driver (Figure 1 and Suppl. Figure S1). Surprisingly, with individual components of the A/B polarity complex containing Baz/Par3, Par6, and aPKC we detected distinct outcomes. While knock-down of aPKC showed classical PCP defects, as expected based on previous work (Djiane et al., 2005) (see also below), both *baz-IR* lines caused wing margin loss in the posterior compartment (Figure 1A-C; Baz protein levels were markedly reduced and barely detectable in the *baz-IR* expressing domain: Figure 1E, and Suppl. Figure S1B-C). The loss of the wing margin is associated with a reduction in Notch signaling or in effects on the Wg signaling pathway, as *wg* is a direct target of Notch signaling along the dorsal/ventral compartment border. Besides the posterior compartment of the wing being slightly reduced, no other patterning defects were observed in the *baz* knockdowns (Figure 1A-C). In contrast to *baz*, knockdown of either *aPKC* or *par6* caused cell lethality with loss of large areas of the posterior wing compartment at 25°C or 29°C (Suppl. Figure S1). When *aPKC-IR* was expressed at reduced levels (driven at 18°C), causing a milder knock-down, the posterior wing compartment was still markedly smaller (Figure 1D; note that wing hairs were also often very short, Fig. 1D-D’, orange arrows), but the wing margin appeared largely normal (Figure 1B’-D’). PCP defects were also observed in the *aPKC-IR* expressing domain (Figure 1D’, green arrow indicating PCP hair orientation), consistent with the previously identified role of aPKC in PCP (Djiane et al., 2005). *Par6-IR* mediated knock-down caused severe loss of wing tissues even at 18°C, making it impossible to correlate these defects with a specific cellular mechanism or signaling pathway (Suppl. Figure S1D-D’). Nevertheless, the wing margin appeared relatively normal in areas where there was no tissue loss (Suppl. Figure S1D-D’, magenta arrows).

**Figure 1.**
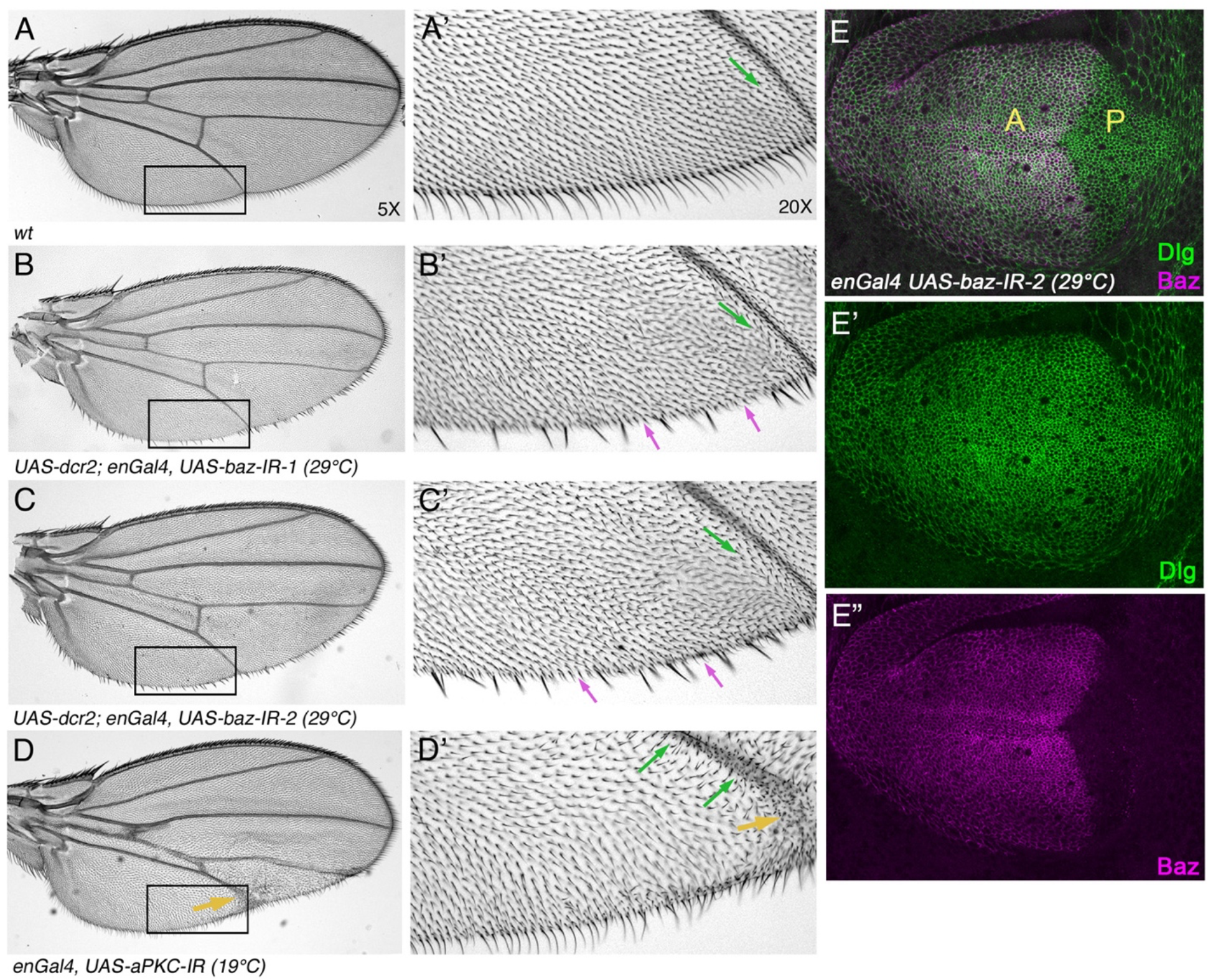
Reduction of *bazooka/baz function* causes wing margin bristle loss. (A-D) Overview and high magnification view (A’-D’) of wings of the indicated genotypes. (A, A’) Wild-type wing; note the associated margin bristles in high magnification of area of posterior wing margin (A’). Cellular hairs point towards the distal margin (green arrow). (B, B’) *enGal4, UAS-baz-IR-1* wing, with *baz* knock-down in the posterior compartment; note loss of margin bristles (magenta arrows). Cellular hair orientation was not significantly affected. (Genotype: *UAS-dcr2; enGal4, UAS-RFP, NRE-GFP / UAS-Baz-IR-1 at 29°C*) (C, C’) *enGal4, UAS-baz-IR2* wing (independent RNAi, see Fig. S1), displaying similar loss of margin bristles phenotype with hair orientation not significantly affected. (Genotype: *UAS-dcr2; enGal4, UAS-RFP, NRE-GFP / UAS-Baz-IR-2 at 29°C*) (D, D’) *enGal4, UAS-aPKC-IR* wing, displaying largely normal margin bristle development. Note wing size reduction in posterior compartment with some areas displaying abnormally short wing hairs (orange arrow) and other areas with cellular hair orientation defects, suggesting that reduction of aPKC affected several aspects of wing development, yet had no effect on margin bristle development (Genotype: *enGal4* /+; *UAS-aPKC-IR (aPKC^JF01966^*) /+ at 19°C) (E-E”) RNAi-based knock down of Baz protein in posterior compartment. All panels are showing max-projections of a stack of confocal images (∼40mm to cover the apical surface of all of the wing pouch area), including composite (E) and individual channels. Anti-Dlg (Discs large, green and E’) staining, serves as positive control to outline cells in both anterior (labeled “A”) and posterior compartments (labeled “P”). Anti-Baz (magenta) levels were markedly reduced in the posterior *baz-IR* expressing compartment, suggesting an effective knock-down via the *baz* RNAi transgene. (Genotype: *dcr2; enGal4/+; /UAS-baz-IR-2 29°C*).

Taken together, knockdown of *baz* caused a distinct loss-of-function (LOF) phenotype from *aPKC* and *Par6*, affecting wing margin development. To confirm this effect, we also analyzed wings containing mutant clones of *baz^4^* or *baz^815-8^* alleles (Suppl. Figure S1A for allele information). Consistent with the RNAi knock-downs, *baz* LOF clones also displayed wing margin loss (Suppl. Figure S2A-C). On the notum, *baz* mutant LOF clones displayed loss of sensory bristles (Suppl. Figure S2D-F), in accordance with previous work (Roegiers et al., 2001) confirming the behavior of the *baz* alleles. Taken together with the RNAi phenotypes above, the *baz* LOF clonal phenotypes in the wing suggest a potential role for Baz/Par3 in Notch-signaling.

### *Notch* and *baz* interact genetically

To corroborate a relationship between *Notch* and *baz* phenotypes, we performed genetic interaction analyses. In heterozygous conditions of the null allele *N^55e11^*/+, approx. 30% of wings displayed distal margin notching (near vein 3 and 4; Figure 2A, B, and F). We compared these wings to double heterozygous combinations with the LOF *baz* alleles: *baz^4^* or *baz^815-8^*. Strikingly, the double heterozygous animals displayed a marked enhancement of wing notching with 68% and 79% of wing margin defects in *N^55e11^*/*baz^4^* and *N^55e11^*/*baz^815-8^* allelic combinations, respectively (Figure 2C-D, and F; see also Suppl. Figure S3). Moreover, in addition to the enhanced distal wing notch phenotype, a novel defect appeared in the double heterozygous animals with wing notching in the proximal region of the posterior compartment (Figure 2B’, D’ and quantified in 2G; see also Suppl. Figure S3B’-C’). Whereas *N^55e11^*/+ wings did not display any margin defects in this proximal region (Figure 2B’ and 2G), 100% of the double heterozygous *N^55e11^*/*baz^4^* and *N^55e11^*/*baz^815-8^*wings carried such margin defects (quantified in Figure 2G; *p<*0.0001). The equivalent *baz^4^*/+ and *baz^815-6^*/+ heterozygous control wings did not display any margin notching phenotypes (Figure 2E-E’, also Suppl. Figure S3C-C’). These data suggest that *Notch* and *baz* genetically interact during wing margin patterning in a synergistic manner.

**Figure 2.**
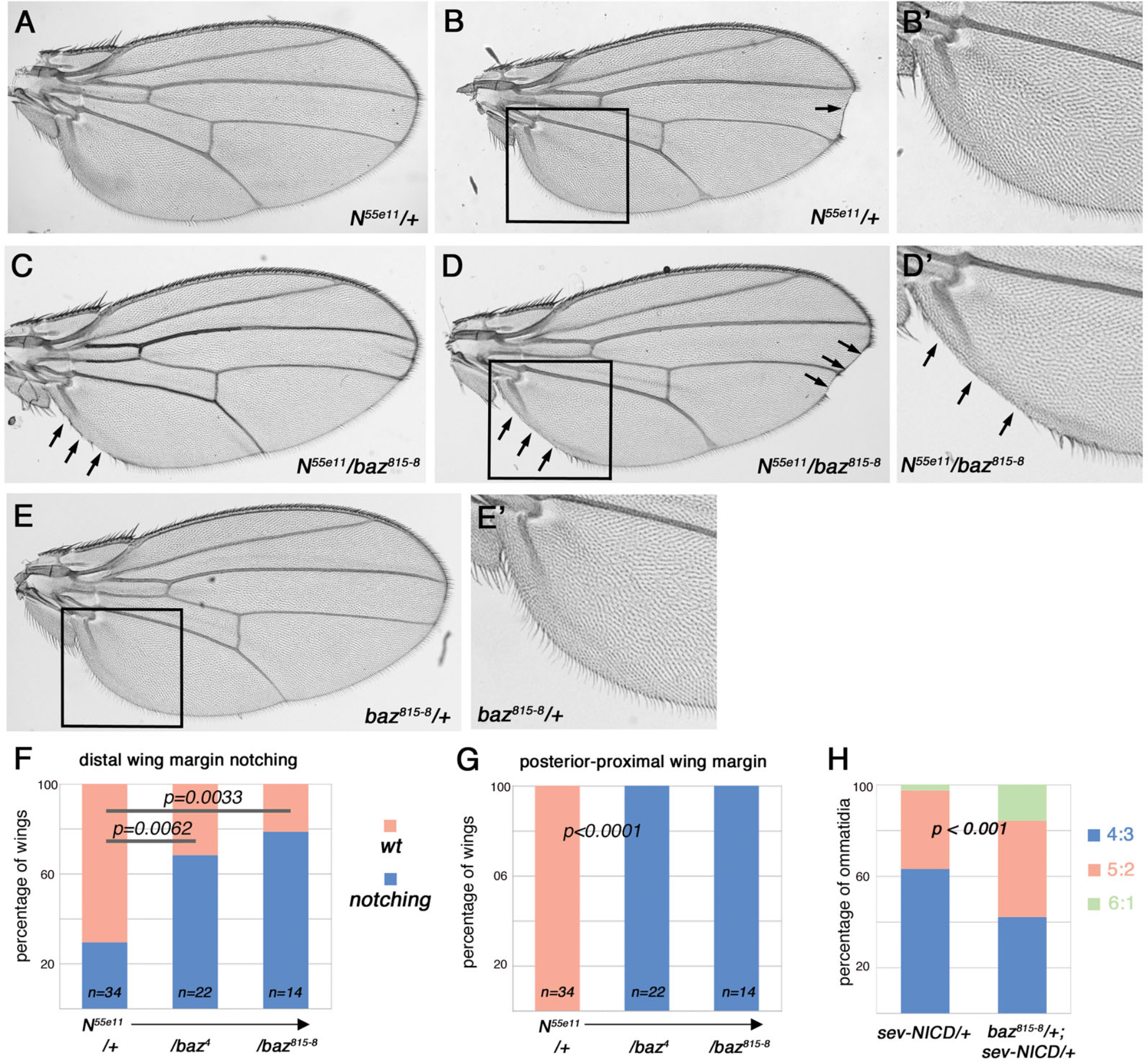
*Notch* and *baz* interact genetically during wing margin patterning. (A-B) Heterozygous *N^55e11^/+* wings display wing margin notching phenotypes; approx. 2/3 of *N/*+ wings are normal with no margin loss (example in A), while 1/3 displayed distal margin notches (B, arrow). Proximal posterior wing margin was always normal (boxed area in B, and B’) in all animals. Quantified in panels (F) and (G). (C-D) *N^55e11^/baz^815-8^* double heterozygous wings. Note increased margin notching in distal area (D, arrows) and proximal-posterior wing margin in all wings (C, D and D’, arrows). (D’) Higher magnification of boxed area in D. Quantified in panels (F) and (G). (E) *baz^815-8^/+* single heterozygous wings show no notching/tissue loss at margin, neither distally nor in the proximal-posterior region (E’, boxed area from E shown at higher magnification). (F) Quantification of distal wing margin phenotypes as percentage of wings containing notching (genotypes as indicated). Note that *N^55e11^/baz^4^* and *N^55e11^/baz^815-8^*flies show enhanced defects (higher percentage) of margin notching as compared to *N^55e11^/+* alone (*p*=0.0062 and *p*=0.0033, respectively (Fisher’s exact test). The control *baz^4^/+* and *baz^815-8^/+* single heterozygous flies have no wing margin phenotypes. (G) Quantification of proximal wing margin phenotypes (posterior compartment) as percentage of wings with margin loss (genotypes as indicated). 100% of *N^55e11^/baz^4^* and *N^55e11^/baz^815-8^*had proximal wing margin notches. *N^55e11^/+, baz^4^/+* and *baz^815-8^/+* heterozygous flies have no proximal wing margin defects. *p*<0.0001 (Fisher’s exact test) for phenotypic differences between both double heterozygous combinations and *N^55e11^/+*. (H) *baz^815-8^/+* heterozygosity suppresses the *sev-NICD*/+ gain-of-function eye phenotype. *sev-NICD* induces several phenotypes, and most prominently a transformation of the R1/R6 outer photoreceptors to R7s, which results in ommatidia classified as 4:3 (4 outer R-cells and 3 R7s) or 5:2 (5 outer R-cells and 2 R7s), in contrast to *wt*, which has a 6:1 appearance (6 outer R-cells and 1 R7). Note that heterozygosity of baz suppresses the effect of sev-NICD and reverts the ommatidial appearance partially towards 6:1 (the *wt* appearance).

We next asked whether interactions between *Notch* and *baz* are also detected in other tissues and with a gain-of-function (GOF) N genotype. The specification of cell fates in the developing eye requires Notch input at multiple steps, from early patterning of the presumptive eye field to the specification of individual photoreceptor (R cell) and accessory cell fates. A classical assay for N-signaling during eye patterning is the expression of constitutively active (ca)-N molecules under control of the *sevenless* (*sev*) enhancer (expressed in R3-4 and R1,6, and R7). Such ca-Notch molecules are either N-terminally truncated molecules that lack the inhibitory region in the extracellular domain preventing g-secretase-mediated cleavage (NDECD, (Fortini et al., 1993)), or the intracellular domain itself, which lacks the membrane targeting sequence (NICD, (Kidd et al., 1998)). Depending upon the strength of activation, ectopic Notch activity induces the transformation of R3 to R4 and/or the R1/R6 photoreceptors to R7 (Cooper and Bray, 1999; Fanto and Mlodzik, 1999; Tomlinson and Struhl, 1999, 2001). The induction of R7 is a classical Notch-dependent cell fate and ectopic activation of N causes the transformation of R1/R6 photoreceptors to R7 independently of any PCP input (Tomlinson and Struhl, 2001). In the *sev-NICD* background a large fraction of ommatidia display a multi-R7 phenotype (Figure 2H, and (Tomlinson and Struhl, 2001)), while the presence of one copy of the strong *baz^815-8^* allele in this background suppresses the GOF Notch effect (Figure 2H). Taken together with the above genetic interactions with *N* LOF features, these results suggest that Baz/Par3 is generally required for Notch signaling function in different tissues and contexts, as seen in both, LOF and GOF, scenarios.

### *baz* loss-of-function leads to reduction of Notch target gene expression

To test whether *baz-IR* affects Notch signaling directly, we examined the expression of the endogenous Notch signaling targets *wg* and *cut* (Micchelli et al., 1997; Neumann and Cohen, 1996). To monitor *wg* expression we analyzed the expression of *wg-lacZ*, an enhancer trap insertion in the *wg* locus, which faithfully reproduces *wg* expression patterns (Kassis et al., 1992); and to monitor *cut* expression we used an antibody to Cut. Knock-down of *baz* in the posterior compartment caused reduction of both *wg-lacZ* expression and Cut protein levels, as compared to control levels in the anterior compartment (Figure 3A-C”). Both *wg-lacZ* and Cut levels were significantly reduced in quantifications of their intensity ratio between the anterior (control) and posterior (experimental) compartment (Figure 3D-E). Since the reduction of *wg* and *cut* expression is detected at the third instar larval stage in wing discs, before sensory development occurs at the pupal stages (Jack et al., 1991), the *baz* effect on their expression precedes sensory cell differentiation and is thus clearly separated in time from the known Baz function in asymmetric SOP cell division.

**Figure 3.**
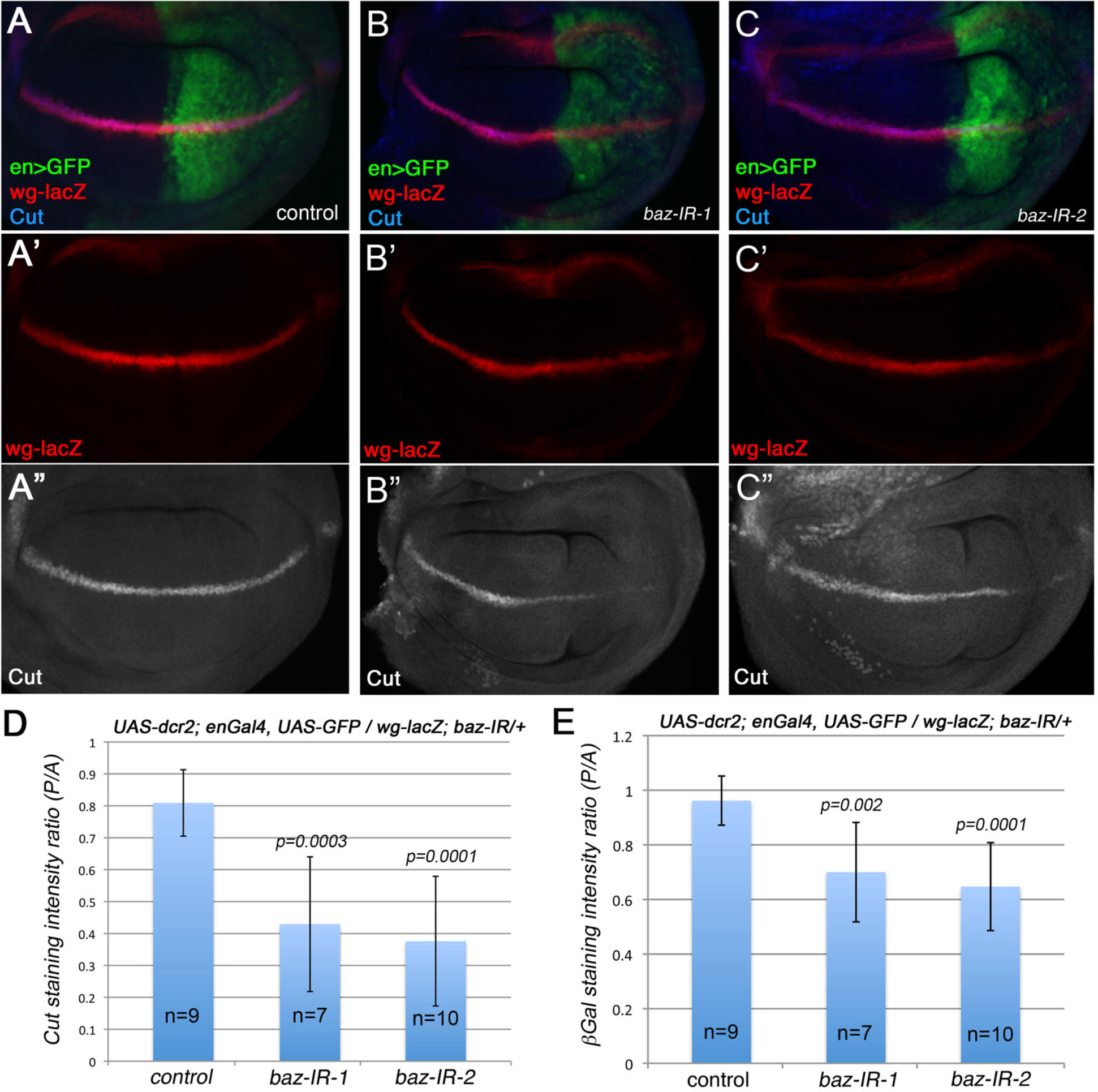
Notch target gene expression is affected in *baz* mutant cells. (A-C”) Confocal images of third instar wing discs with expression of Notch target genes: *wg-lacZ* (detected with anti-bGal, red; monochrome in A’, B’, and C’) and *cut* (anti-Cut, blue; monochrome in A”, B”, and C”) are shown in wild type (A-A”), *baz-IR-1* (B-B”), and *baz-IR-2* (C-C”) genotypes. *wg-lacZ* and Cut expression are both reduced within the *enGal4*>*IR* expression domain (posterior compartment, marked by GFP in green). Genotypes: (A) *UAS-dcr2; enGal4, UAS-GFP / wg-lacZ*. (B) *UAS-dcr2; enGal4, UAS-GFP / wg-lacZ; baz-IR-1* (*BL31522)/+*. (C) *UAS-dcr2; enGal4, UAS-GFP / baz-IR-2 (BL31523)/+* (all at 29°C). (D) Quantification of intensity ratio of Cut expression between anterior and posterior compartments (see Materials and Methods for details). Note reduction with both independent *baz-IR* transgenes (*p*-values 0.0003 and 0.0001 (student t-test), respectively). (E) Quantification of intensity ratio of *wg-lacZ* expression (anti-bgal staining) between anterior and posterior compartments (*p*-values 0.002, and 0.001, respectively).

To confirm these results, we also examined the expression of a synthetic reporter containing the Notch response element (NRE), *NRE-GFP* (Furriols and Bray, 2001; Saj et al., 2010). In wing discs, *NRE-GFP* is expressed along the dorsal/ventral compartment border in pattern indistinguishable from endogenous Notch target genes such as *wg* and *cut* (Saj et al., 2010) (Suppl. Figure S4A). Importantly, reduced expression of *NRE-GFP* was observed in the posterior compartment in *enGal4>baz-IR(s)* wings (Suppl. Figure S4B-C’, and E), indicating that a reduction of *baz* directly correlated with reduced Notch-signaling and target gene expression.

To corroborate these data and to test for a general effect of *baz* LOF on Notch signaling outcomes in different experimental set ups, we analyzed the expression of Notch targets in *baz* mutant clones. Consistent with the *baz-IR* knockdown in wing discs, Cut expression levels were reduced in *baz^815-8^* mutant cells (clones were positively labeled by RFP expression, Figure 4A-C, examples highlighted with white arrows; quantified in Figure 4D). Next we tested for effects on Notch signaling in eye discs. To test an independent Notch target gene, we asked whether expression of *atonal* (*ato*), a N-signaling target during early eye patterning prior to R-cell specification (Freeman, 2001) also requires *baz* for its full expression. Ato levels were reduced in *baz^815-8^* mutant cells (orange arrows in Figure 4E-E”, F-F”, mutant clone marked by lack of GFP). Importantly, apical-basal polarity as apparent in optical z-sections was normal in *baz*^-^ clones (Figure 4G-G”; note Patj as A/B marker being localized normally in *baz^815-8^* clones – see Discussion), whereas a reduction of Ato expression was also easily detected in this optical plane (Figure 4D-F, orange arrows in E” and F”; quantified in Figure 4H),

**Figure 4.**
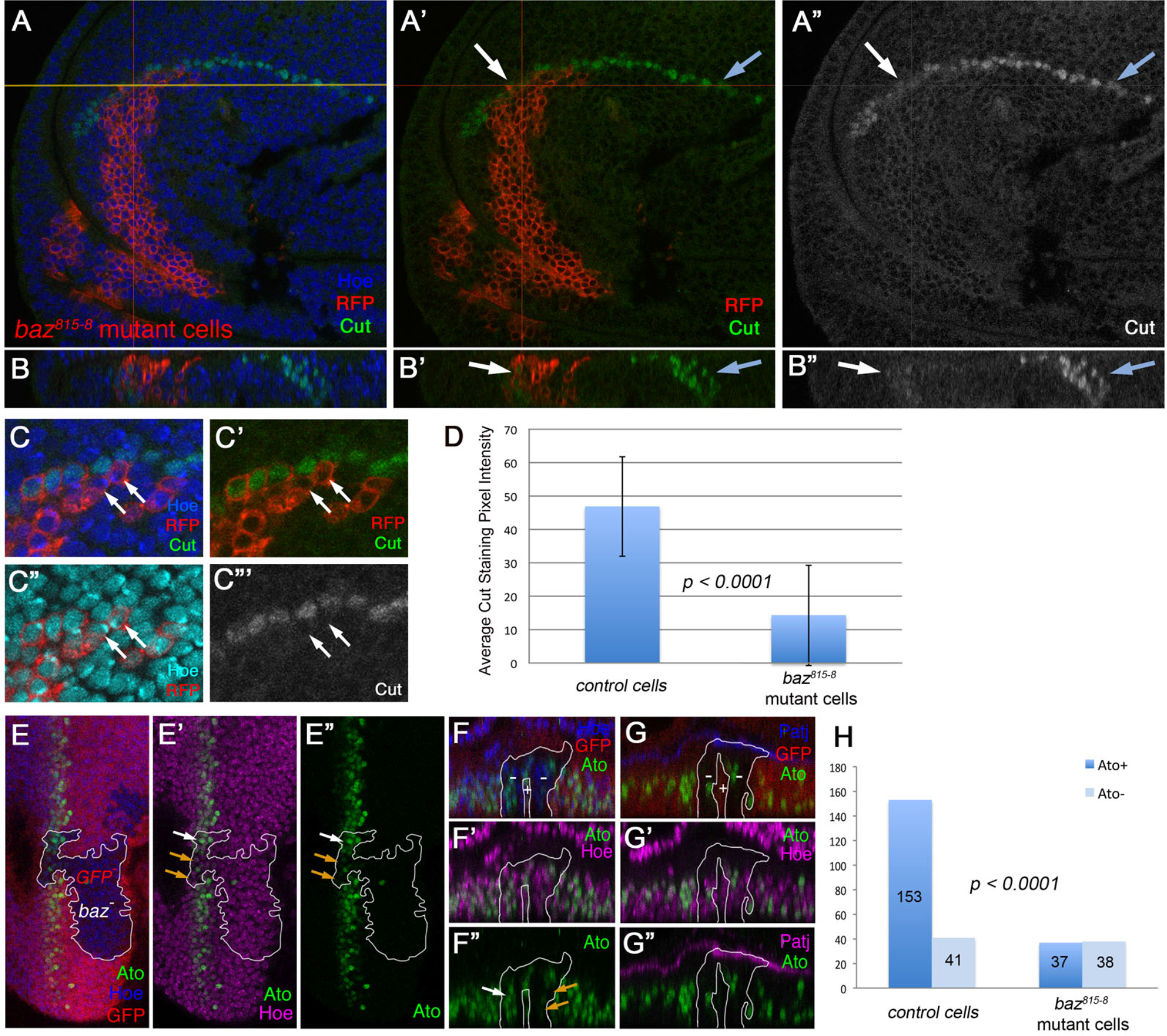
Clones mutant for *baz* display markedly reduced Notch target gene expression in eye and wing discs. (A-A”) Confocal optical section of *baz^815-8^*wing disc clone (positively labeled by co-expression of RFP), and (B-B”) cross-section of same *baz*^815-8^ clone at position indicated by yellow line in (A). Cut expression is reduced in *baz* mutant cells. Yellow line traverses Cut expression domain in two regions indicated by white and blue arrows. Cut levels in *baz*^815-8^ mutant cells (white arrow) are reduced, as compared to heterozygous control cells (blue arrows). Genotype: *w, tub-gal80, hs-flp, FRT19A/ baz^815-8^ FRT19A; act-Gal4, UAS-mCD8RFP /+.* (C-C”’) Projection of 3 optical sections at clonal region in D/V border cells, showing Cut (green, monochrome in C”’) expressing cells. Cut is normally expressed in two rows, each flanking the D/V-boundary on the respective side. Note that *baz^815-8^* mutant cells (RFP+, in red) in the cells abutting the D/V-boundary express Cut at markedly reduced levels or not at all (examples highlighted by white arrows), nuclei are marked by Hoechst (blue in C, or turquoise in C”). (D) Quantification of Cut expression levels of D/V border cells (measured by average pixel intensity for each cell, see Materials and Methods for details), plotted to compare Cut levels between control and *baz*^815-8^ mutant cells. Note higher Cut expression in neighboring wild-type cells, as compared to *baz*^815-8^ mutant cells. (E-E”) Atonal (Ato) expression at and near the morphogenetic furrow (MF) in eye imaginal discs. containing *baz*^815-8^ mutant clone (marked by absence of GFP expression, red, highlighted by white outlines). Note markedly reduced Ato expression (green) in many mutant cells (orange arrows in E’, E”), although some remained unaffected (E’ and E”, white arrow). All cells, including those with low Ato expression levels, stained equally well with nuclear marker (Hoechst, blue in E, magenta in E’). (F-G”) Two z-section views of *baz*^815-8^ mutant clone in furrow region, with (F) scanned through the middle of Ato expression stripe. Note that many *baz*^815-8^ mutant cells displayed markedly reduced or lost Ato expression (orange arrows in F”) with Hoechst (Hoechst, blue in F,G and magenta in F’,G’) staining serving as control. (G) z-scan through posterior region of Ato expression stripe, where scan-plane intersected with the apical domain (due to tissue shape of the MF and pseudo-stratified epithelium of imaginal disc) reflecting junctional marker Patj (blue in G, magenta in G”). Note that apical junctional region remains intact in *baz^815-8^* mutant cells, indicating that apical-basal polarity was not affected. Genotype: *baz^815-8^ FRT9-2/hsflp ubi-GFP FRT9-2*. (H) Quantification of Ato expressing cells in *baz^815-^* clones and neighboring wild-type control cells revealed a decrease in Ato positive cells in *baz^815-^*mutant clone vs wild-type control. Distributions were significantly different based on Fisher’s exact test.

Taken together, these results are consistent with the notion that Baz functions as a positive regulator within the Notch-signaling pathway, promoting the expression of Notch target genes.

### Baz affects Notch-signaling at the NICD level

To mechanistically dissect Baz function in Notch signaling, we used N-terminal Notch truncations in GOF assays to determine at what level within the pathway Baz was required. First, we used NDECD (Fortini et al., 1993), which lacks the extracellular domain and is constitutively active independent of ligand binding (see above). As such, if NDECD were affected by *baz* LOF, Baz would be required downstream of the N-ligands and the subsequent extracellular cleavages of Notch, but upstream of the g-secretase-mediated cleavage (Andersson et al., 2011; Struhl and Greenwald, 1999). When NDECD was clonally expressed in larval wing discs, 43% of NDECD clones induced the Notch target *wg* (Figure 5A, F; as detected by Wg antibody). When *baz-IR(*s) were co-expressed with NDECD, the frequency of clones expressing Wg was reduced to 20% (*baz-IR*-1) and 27% (*baz-IR*-2), respectively (Figure 5B, F). These data suggested that Baz was required at the level of the g-secretase cleavage or further downstream within the Notch-signaling pathway. Consistent with this hypothesis *baz-IR* knock-down displayed no detectable effect on Notch protein levels at the membrane or cellular distribution of Notch (Suppl. Figure S1E-F).

**Figure 5.**
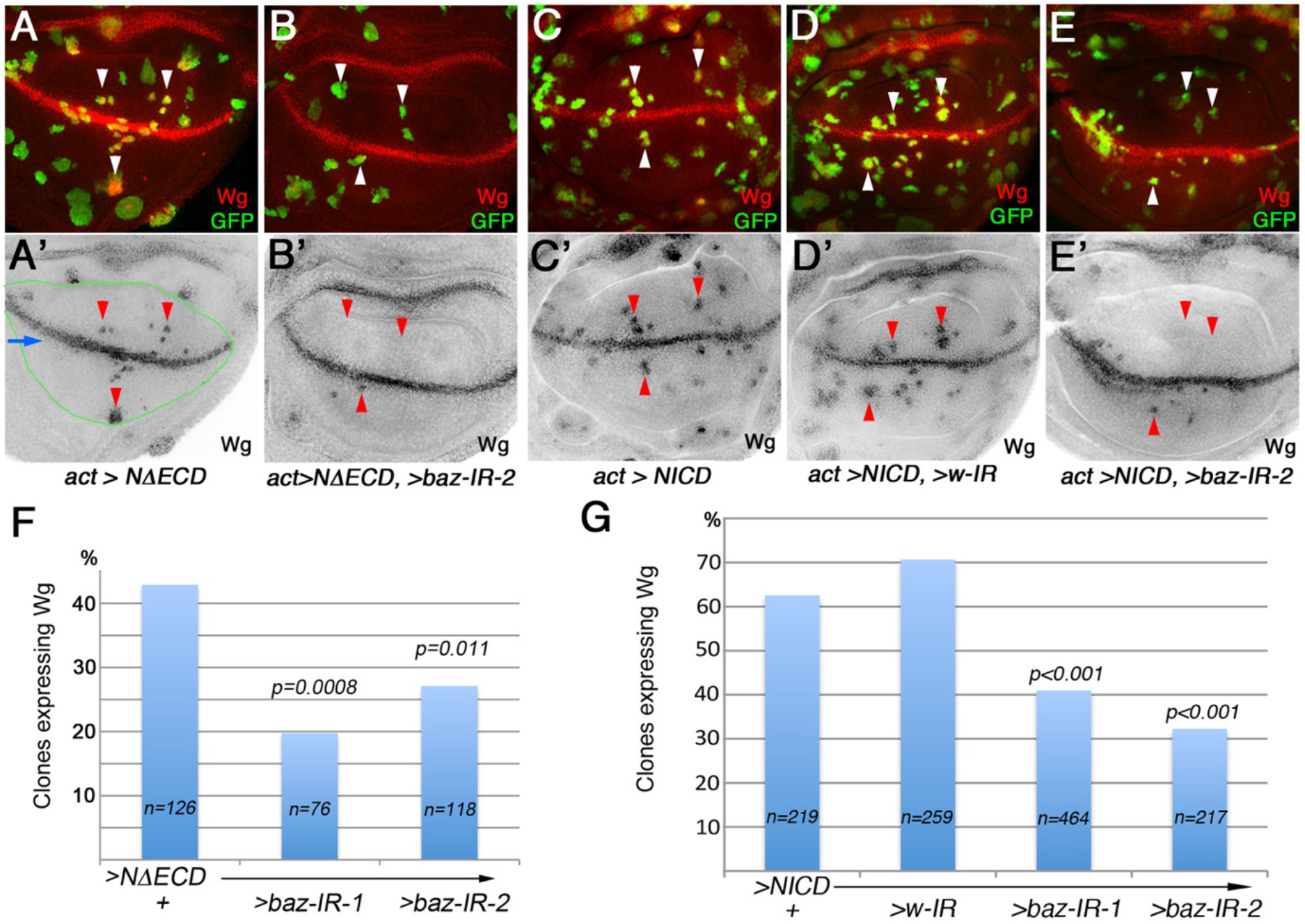
Baz acts at the level of NICD or further downstream within Notch signaling. (A-A’) NDECD expressed via flip-out clones induces ectopic Wg expression within wing blade precursor cells (Wg in red, monochrome in A’; clones labeled by co-expression of GFP in A). Endogenous Wg expression stripe at DV-boundary serves as control (blue arrow in A’) here and in other genotypes, clones overlapping with endogenous Wg stripe were not scored in quantification. Arrowheads mark examples of NDECD expressing clones (white in A, red in A’), many of which induce Wg expression. Only clones in the wing pouch proper (pouch outlined by green lines) were scored for Wg expression. Genotype: *hsflp/UAS-dcr2; act>y^+^>Gal4, UAS-GFP/+; UAS-NDECD /+*. (B-B’) Clonal co-expression of NDECD with *baz-IR-2* markedly reduced the number of Wg expressing clones (Wg in red, monochrome in B’; clones labeled by co-expression of GFP). Arrowheads mark examples of NDECD expressing clones (white in B, red in B’); note reduction in such clones expressing Wg (compare to A,A’) Genotype: *hsflp/ UAS-dcr2; act5c>y^+^>Gal4, UAS-GFP/+; UAS-NDECD /baz-IR-2*. (C-C’) NICD expressing clones induce ectopic Wg expression with high frequency (Wg in red, monochrome in C’; clones labeled by GFP). Arrowheads highlight examples of NICD expressing clones (white in C, red in C’). Genotype: *hsflp/UAS-dcr2: act5c>y^+^>Gal4, UAS-GFP/+; UAS-NICD /+*. (D-D’) *IR* control (*white*/*w-IR*) co-expressed in NICD in clones. Ectopic Wg expression was induced at similar frequencies to NICD alone (Wg in red, monochrome in D’; clones labeled by GFP). Arrowheads highlight examples of NICD expressing clones (white in D, red in D’). Genotype: *hsflp/UAS-dcr2: act5c>y^+^>Gal4, UAS-GFP/ UAS-NICD /+*; *w-IR/+*. (E-E’) Clonal co-expression of NICD with *baz-IR-2* markedly reduced the number of Wg expressing clones (Wg in red, monochrome in E’; clones labeled by GFP). Arrowheads highlight examples of NICD expressing clones (white in E, red in E’). Genotype: *hsflp/ UAS-dcr2; act5c>y^+^>Gal4, UAS-GFP/UAS-NICD; baz-IR-2/+*. (F) Quantification of NDECD clones with induced Wg expression with or without *baz-IR(s)*. The frequency of Wg activation in >*NDECD* clones was markedly reduced by the presence of *baz-IR-1 or baz-IR-2* (*p*=0.0008 or *p*=0.011, respectively). Only clones inside wing pouch region were counted, due to *wg* expression being regulated by other pathways outside pouch region. (G) Quantification of NICD expressing clones with induced Wg expression with genotypes as indicated (with *w-IR* (control), *baz-IR-1*, and *baz-IR-2*). Wg expression frequency in NICD clones was markedly reduced when *baz-IR-1 or baz-IR-2* were co-expressed (both *p*<0.001 comparing to NICD alone or NICD with *w-IR*; as calculated by Fisher’s exact test).

To further investigate the position of Baz within the Notch pathway, we tested the NICD fragment, which is fully active and does not associate with the membrane or require any cleavage and is thought to directly translocate to the nucleus to activate Notch-signaling target genes as transcriptional co-activator (Andersson et al., 2011; Bray, 2016; Struhl and Greenwald, 1999). In control wing disc clones expressing NICD, Wg expression was readily detected in 63% of clones (or 71% with a control IR co-expression) (Figure 5C-D, and G). In experimental clones, with *baz-IR*(s) being co-expressed with NICD, the NICD effect was markedly suppressed: to 41% with *baz-IR*-1 and 32% with *baz-IR*-2, respectively (Figure 5E, G). These data are consistent with the notion that Baz acts downstream of membrane tethered Notch and that it is required for the NICD fragment to activate Notch-signaling target gene expression.

In the nucleus, NICD functions as a co-activator interacting with Su(H) (the *Drosophila* member of the CSL transcription factor family), Mastermind (Mam), and other co-activators to form a complex (or complexes) at transcriptional enhancer sites to activate Notch target gene expression (Bray, 2016). To gain insight into whether Baz/Par3 promotes the formation of the Su(H)-NICD-Mam complex, we asked *whether baz* LOF could reduce the activity of an active Su(H)-Ank fusion protein. A direct fusion of the Ankyrin repeats of Notch to Su(H), was shown in *Xenopus* to bypass a requirement for endogenous Notch itself by directly recruiting Mam, and thus it is acting even further “downstream” relative to NICD in the pathway (Wettstein et al., 1997). To this end, we generated a *Drosophila* version of the construct, fusing the Notch ankyrin repeat region to the Su(H) protein [yielding Su(H)-Ank]. We used the clonal expression assay in the wing pouch, as used above for NICD for example (Figure 5), and expressed Su(H)-Ank in equivalent clones with *baz-RNAi*. Strikingly, clonal expression of Su(H)-Ank alone or together with *baz-RNAi* showed similar percentages of clones inducing expression of *wg* (Suppl. Figure S5). This result demonstrates that Su(H)-Ank activates Notch target gene expression independently of Baz. Taken together with the clonal behavior of NICD, these data suggested that Baz acted on NICD, upstream of the Su(H) based transcription complex, prior to NICD association with Su(H) and Mam (see Discussion).

To complelement the above *in vivo* studies we next addressed whether Baz and NICD could physically interact. This was tested biochemically via co-immunoprecipitation (co-IP) in S2 cells that were co-transfected with HA-tagged NICD and either Baz-myc (full length Baz tagged with myc) or myc-Hippo (Hpo, as negative control). Protein extracts were immunoprecipitated with anti-myc antibody and analyzed by immunoblotting with anti-HA antibody. NICD was readily co-immunoprecipitated with Baz-myc (Figure 6A), but not by the Hpo negative control (Figure 6A-B; see figure legend for additional controls), suggesting that NICD and Baz/Par3 can physically associate.

**Figure 6.**
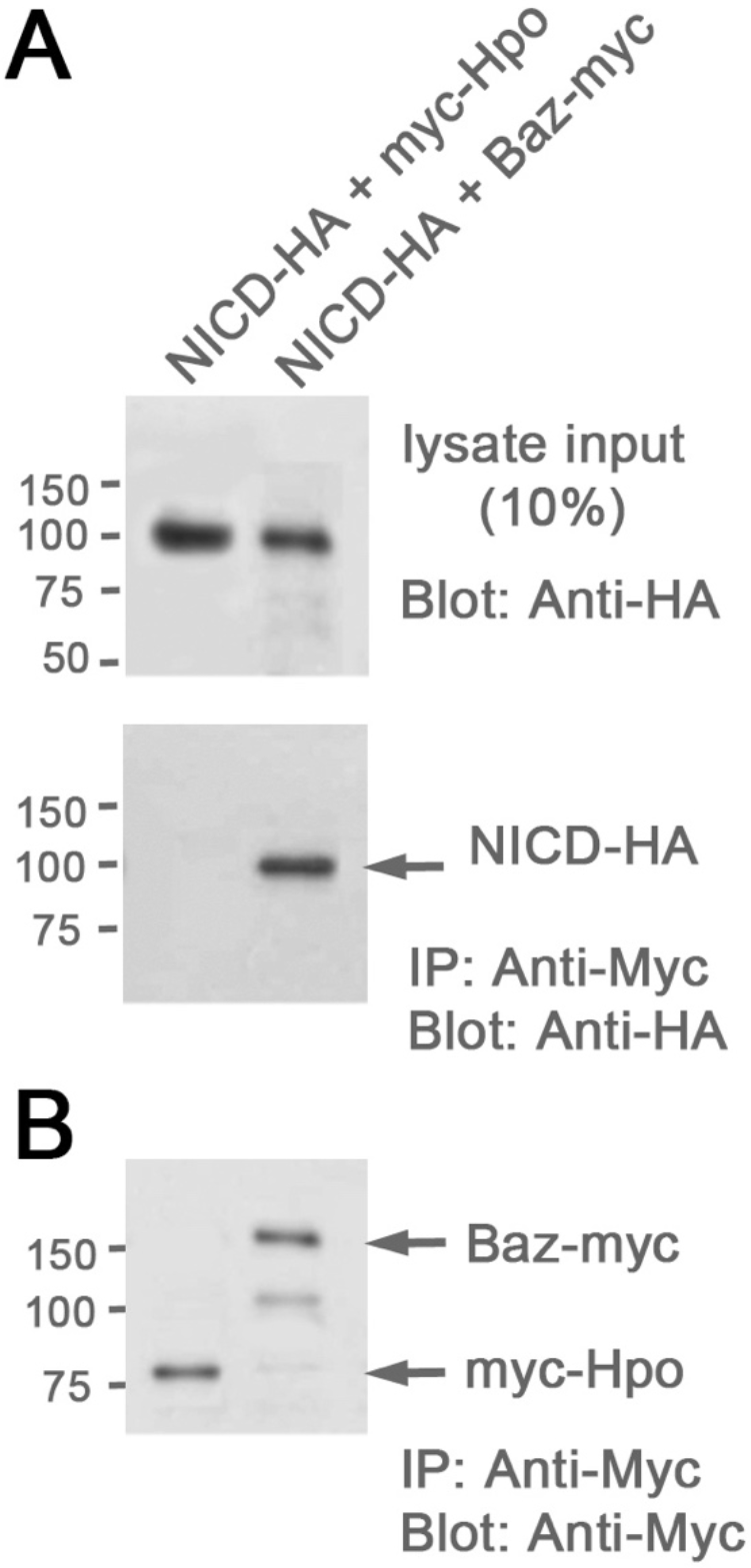
Baz is detected in the same protein complex as NICD. Baz (myc-tagged) or myc-Hippo (Hpo, negative control) and HA-tagged NICD were co-transfected into S2 cells. Lysates of transfected cells were immnunoprecipitated (IP-ed) with anti-myc antibody. (A) Upper panel: Western blot showing NICD-HA detected in lysates of cells co-transfected with myc-Hpo or Baz-myc. Lower panel: Anti-myc co-IP experiments, note NICD-HA was co-IPed with Baz-myc, whereas it was not IP-ed by myc-Hpo (specificity control). (B) Both Baz-myc and myc-Hpo were efficiently pulled down with anti-Myc in cell lysates co-transfected with NICD-HA.

Taken together with the functional *in vivo* data, these results suggested that Baz and NICD were associated in the same protein complex, and that the presence of Baz is required for normal NICD function in Notch target gene induction. However, Baz appeared to act upstream of the Su(H) based transcription factor complex, as it did not have an effect in the context of the Su(H)-Ank fusion, which bypassed a requirement for NICD.

### Baz and Numb are required at different levels within the Notch pathway

To gain further mechanistic insight into how Baz might affect NICD, we tested the relationship of Baz to Numb, which has itself a complex relationship to Notch-signaling (see also Introduction). While Numb has been shown to regulate Notch-signaling at the level of trafficking of the receptor (Couturier et al., 2013; Houssin et al., 2021; Schweisguth, 2015), it has also been reported to directly interact with NICD and inhibit Notch signaling at the NICD level (Frise et al., 1996; Guo et al., 1996). To address these possibilities, we wished to test whether *numb* and *baz* could interact genetically with each other in the above wing margin specification context. First we asked, whether reduction of *baz* function (via *baz-RNAi* equivalent to experiments shown in Figure 1) could be modified by Numb levels. Indeed, loss of wing margin bristle cells (Figure 7A, C, D, F) was suppressed upon removal of one *numb* gene copy, in a *numb^15^/+* heterozygous background (Figure 7B, C, E, F), displaying a dosage sensitive relationship between the two proteins. To confirm this, we compared phenotypes induced by overexpression of *numb-GFP* with or without *baz-Flag* in developing wings. Overexpression of *numb-GFP* suppressed Notch signaling, as detected by margin bristle loss (Suppl. Figure S6A-B and D, (Frise et al., 1996); of note, Numb-GFP also inhibited Notch function in the context of wing vein formation). Co-overexpression of *baz-Flag* with *numb-GFP* reduced the inhibitory effect of Numb on Notch signaling at the wing margin (Suppl. Figure S6B-D). These experiments support the notion that *numb* and *baz* antagonize each other genetically during wing margin cell induction with Baz promoting Notch function and Numb inhibiting it.

**Figure 7.**
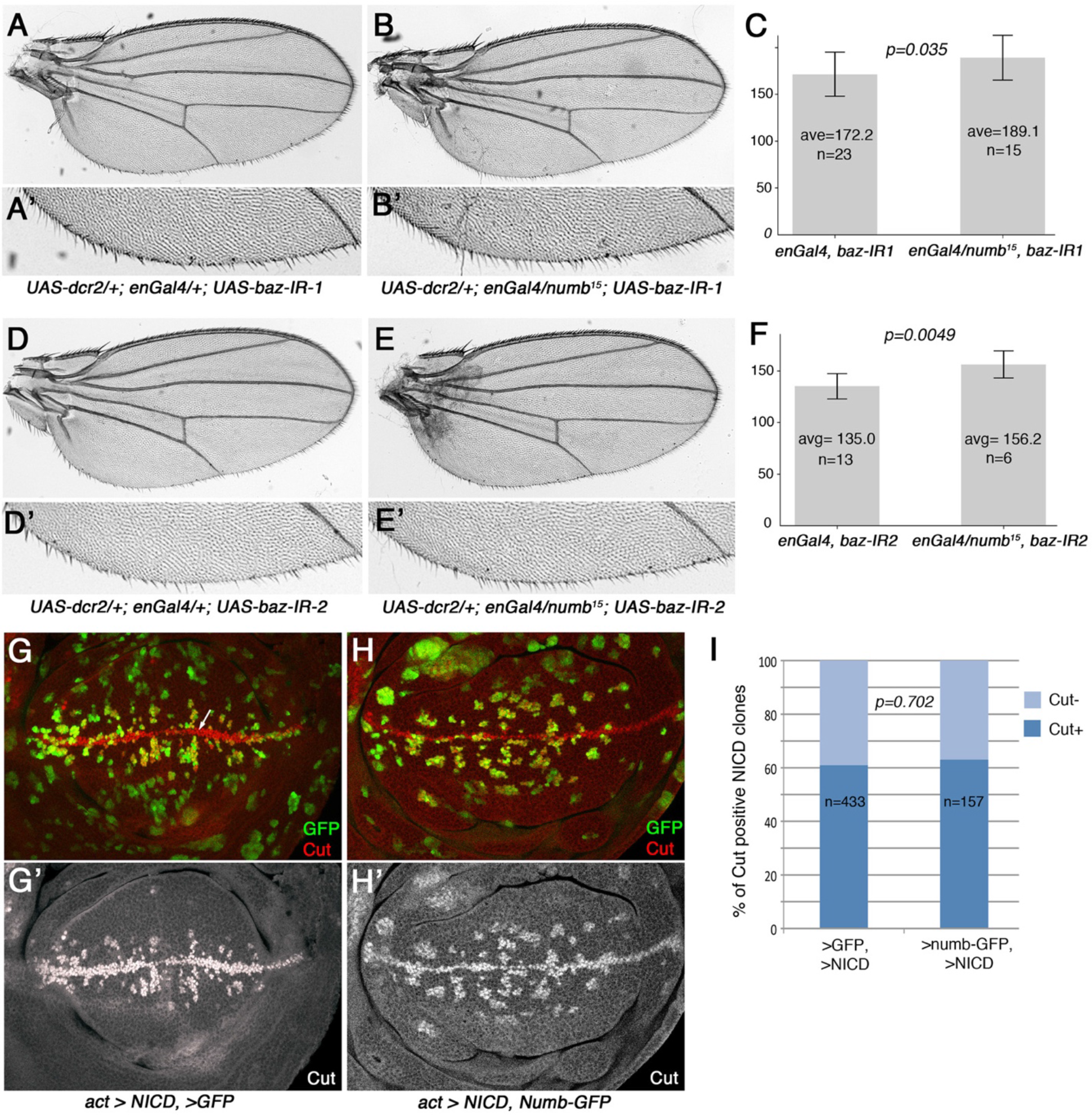
*baz* and *numb* act antagonistically to each other, but at different levels within the Notch pathway. (A-F) Wings expressing *baz RNAi* in the posterior compartment in wild-type (*baz-IR-1* in A,A’ and *baz-IR-2* in D,D’) causing wing margin bristle cell loss (also compare to Figure 1) or the equivalent *baz-IRs* in a *numb* heterozygous background, *numb^15^/+* (in B,B’ and E,E’, respectively). Quantifications, as bristle number, in (C) and (F); wild-type posterior wing margin has on average 230 bristle cells (Suppl. Figure S6). Note that removing a copy of *numb* suppresses the effect of either *baz-RNAi*. Experiments were carried out at 29°C. Genotypes: (A, A’) *UAS-dcr2/+; enGal4/+; baz-IR-1/+*. (B,B’) *UAS-dcr2/+; enGal4/numb^15^; baz-IR-1/+,.* (D,D’) *UAS-dcr2/+; enGal4/+; baz-IR-2/+*. (E,E’’) *UAS-dcr2/+; enGal4/ numb^15^; baz-IR-2/+,.* Note that *baz-IR-2* causes stronger defects, and that the phenotypic suppression caused by *numb^15^ heterozygosity* was significant (t-test) in both cases (C and F, respectively). “n” = number of wings analyzed. A’, B’, D’, and E’ show higher magnification of the corresponding wings shown. (G-I) Induction of Notch target gene expression by NICD expressing clones was not affected by Numb co-overexpression. NICD was expressed via flip-on clones, as in Figure 5, with clones labeled with GFP or numb-GFP co-expression. (G) Confocal optical projection with NICD and GFP expressing clones (control). Note ectopic induction of the Notch target gene *cut*, visualized via anti-Cut immunohistochemical staining (red and monochrome in G’), in many NICD clones. Cut is normally expressed in a stripe at the dorsal/ventral boundary (arrow, in panel G). Genotype: *hsFlp/+; act5c>y>Gal4, UAS-GFP/UAS-NICD*. (H) Projection of wing disc confocal optical sections containing NICD expressing clones (marked in green by Numb-GFP co-expression) with Numb-GFP overexpressed together with NICD. Note ectopic induction of *cut* expression in many clones within the wing pouch. Genotype: *hsFlp/+; act5c>y>Gal4, UAS-numb-GFP/UAS-NICD*. (I) Quantification of the percent of Cut expression positive clones in NICD/GFP and NICD/numb-GFP expressing clones. Note similar ratio of expressing clones in both genotypes with no significant difference (*p* = 0.702 based on Fisher’s exact test; “n” indicates the number of clones).

We next wished to test whether they could act at the same level in Notch-signaling, at the level of NICD. To this end, we asked whether the NICD clonal expression phenotype, induction of target gene expression, which is suppressed by *baz-IR* (see Figure 5 in Results above), could also be affected by Numb levels. Again, clonal NICD expression in the wing pouch induced Notch-signaling target genes, for example *wg* (Figure 5) and *cut* (Figure 7G-G’) and the removal of *baz* (via RNAi) in such clones reduced the ability of NICD to induce target gene expression (see Figure 5E,G). We thus asked whether co-overexpression of *numb-GFP* in such NICD expression clones could also suppress NICD’s function in target gene activation. However, we did not observe an effect of Numb-GFP overexpression on NICD function in target gene expression (Figure 7G-H, quantified in 7I, as shown for Cut). Taken together with the genetic interactions above, these data suggested that Numb and Baz act at different levels in the Notch pathway, with Numb acting further upstream and Baz/Par3 acting at the level of NICD.

## Discussion

In this study we define a novel, direct, and positive role of the polarity protein Baz/Par3 in the Notch pathway. This stands in contrast to its previously established role in Notch signaling regulation during asymmetric SOP cell divisions, where Baz/Par3 affects N-signaling indirectly by asymmetrically preventing the inheritance of the Notch inhibitor Numb (Schweisguth, 2015). In the SOP scenario, the Baz/Par3-Par6-aPKC protein complex localizes to one side within the dividing cell, and a Numb-Neuralized containing complex localizes to the other side of the cell. As Numb inhibits Notch signaling by regulating Notch-Sanpodo trafficking and thus reducing Notch levels at the cell surface, this asymmetry causes Notch inhibition in the Numb containing daughter cell (reviewed in (Schweisguth, 2015)). In the SOP context, Baz/Par3 has been thus proposed to have a passive role in N-signaling activation, via inhibiting an inhibitor, Numb. Our data are consistent with the notion that Baz/Par3 is also required positively for the Notch signaling response and full transcriptional activation of Notch targets in *Drosophila* in multiple cell types. Genetic epistasis studies indicate that Baz acts at the level of cleaved active Notch, NICD, or in parallel to it. Consistent with this notion Baz and NICD co-immunoprecipitated, suggesting that Baz promotes Notch signaling at the level of NICD.

### A novel Baz/Par3 function independent of the A/B polarity complex

Baz/Par3 is generally associated with Par6-aPKC at early to intermediate stages of A/B polarity establishment, but later in mature epithelia it is no longer essential for their maintenance (Morais-de-Sa et al., 2010; St Johnston and Sanson, 2011; Tepass, 2012). Indeed, the fact that we can recover *baz* LOF clones at all, whereas Par6 and aPKC null alleles are cell lethal, suggests that *baz* can behave differently to the ‘core’ A/B factors. In our experimental set-up we observed no evidence for a Par6 or aPKC involvement in Notch signaling. Our data suggest that Baz/Par3 regulates NICD independently of its role in the A/B polarity complex. Similar to A/B polarity establishment, in the context of planar cell polarity (PCP) patterning, Baz/Par3 functions together with aPKC to regulate Frizzled levels (and hence the PCP outcomes) in the *Drosophila* eye (Djiane et al., 2005), and thus, while Par6 could not be assayed in this context, these data suggested that in PCP determination Baz is functioning with established A/B polarity factors. As such, the discovery that in Notch signaling Baz/Par3 appears to function independently of its partners in A/B establishment within the same tissue is unexpected.

Par3 homologues have however been shown to act independently of the aPKC-Par6 complex in several scenarios in mammalian cells, including Hippo/Yap signaling and during epithelial-mesenchymal transition (Zhang et al., 2016). Par3 has even been suggested to act with Ku70 and Ku80 as part of the DNA-dependent protein kinase complex to regulate double-strand breaks (Fang et al., 2007).

### Why was Baz/Par3 not identified earlier as a component of Notch signaling?

The Notch pathway genes were initially identified through classic genetic collections of spontaneous mutants showing morphological defects in *Drosophila* and *C. elegans* (Fleming et al., 1990; Greenwald, 1985; Kopczynski et al., 1988; Lindsley and Zimm, 1992). Subsequent genetic screens and molecular studies have added many components of the Notch pathway and many regulatory mechanisms that affect Notch signaling at all levels, ranging from receptor and ligand trafficking and membrane stability and its membrane associated proteolytic cleavages to nuclear co-factors and chromatin associated target gene regulation (reviewed in (Bray, 2016; Henrique and Schweisguth, 2019; Kopan and Ilagan, 2009). In particular, many of the screens were performed using the notum Notch associated phenotypes and score for asymmetric division-dependent SOP/bristle patterning defects. It has thus been technically difficult, if not impossible, to separate potential additional Baz/Par3 involvements in Notch signaling. Similarly, classical loss-of-function screens in the *Drosophila* embryo would have missed Baz/Par3 due to its initial requirement in epithelial polarization that could have caused earlier lethality (Harris and Peifer, 2004; Muller and Wieschaus, 1996), which obscures a potential role in subsequent signaling events. Indirectly through our screen asking whether A/B polarity factors can also play a role in PCP establishment in the posterior wing disc compartment we used a unique experimental scenario: first the margin loss is due to tissue loss and not asymmetric division of innervated SOPs, and, second, at this later stage of the epithelial sheet is fully formed and Baz/Par3 is not required for maintenance of A/B polarity. This combination thus allowed the identification of *baz* as a novel factor that positively regulates Notch signaling in general.

### Baz/Par3 has multiple independent functions during Notch signaling

Baz/Par3 and Numb have both been implicated in the regulation of Notch signaling via trafficking of the Notch receptor (Couturier et al., 2013; Houssin et al., 2021; Schweisguth, 2015). In addition, independent of trafficking, Numb has been suggested to inhibit Notch signaling directly by binding to NICD (Frise et al., 1996; Guo et al., 1996). Overexpression of Numb indeed can interfere with Notch signaling at the wing margin, causing significant Notch loss-of-function type defects ((Frise et al., 1996) and this work). To address this directly, we asked whether Numb overexpression could interfere with NICD function in the context Notch target gene activation. However, NICD activity was not affected by Numb overexpression, suggesting that it acted further upstream, likely as suggested at the level of regulating Notch trafficking, which is consistent with previous studies (Couturier et al., 2013; Houssin et al., 2021). In contrast to *numb*, knockdown of *baz/par3* directly affected NICD function and reduction of Baz/Par3 levels significantly reduced NICD function in target gene activation. Therefore, Baz/Par3 can act at two independent levels within Notch signaling, affecting (i) membrane trafficking of the Notch receptor (Couturier et al., 2013; Houssin et al., 2021; Schweisguth, 2015) and (ii) serving a positive function downstream in the pathway at the NICD level (see below).

### Baz acts at the NICD level affecting expression of N-signaling targets positively

The Notch-signaling associated DNA binding complex is centered around the Notch-pathway dedicated DNA-binding factor CSL, Su(H) in *Drosophila*, and it switches from a repressor complex to a transcriptional activator complex via NICD (Bray, 2016). NICD binds to Su(H), and recruits co-activators including Mastermind (Mam), and its binding to Su(H)/CSL displaces the “repressor complex” and induces N-signaling target gene transcription. Several factors have been identified that associate with Su(H)/CSL and NICD (see reviews (Borggrefe and Oswald, 2009; Bray, 2016)). Genetic epistasis data, both in the wing and the eye, position the function of Baz/Par3 at the level of NICD after its last cleavage, suggesting that Baz might act in the nucleus together with NICD or alternatively it might promote stability, nuclear translocation, function, or retention of NICD. The biochemical immunoprecipitation results support the genetic data and are consistent with the notion that Baz is a direct regulator of Notch signaling, acting at the level of NICD. However, the Su(H)-Ank fusion appears to not require Baz/Par3 for its capability to activate Notch target gene expression, with *baz* knockdown having no effect on Su(H)-Ank mediated activation, unlike with NICD, suggesting that Baz is acting upstream of the Su(H)-Ank association. The Ank repeats of NICD are critical within the Notch protein for its nuclear response as part of the Su(H) transcription factor complex by recruiting Mastermind, and as such a Su(H)-Ank fusion protein is sufficient to activate Notch signaling target gene expression (Collu et al., 2012; Wettstein et al., 1997). As *baz* mutations affect the levels of target gene expression and the Baz protein acts genetically at the level of, or in parallel to, NICD, Baz/Par3 might be a factor participating in regulating NICD localization, stability, or modifications to mediate its association with Su(H).

Interestingly, the human Baz/Par3 homologue, PARD3, has been suggested to act at a similar level in the YAP/TAZ pathway. PARD3, which itself is found in the nucleus in addition to the junction-bound aPKC/Par6 complex, has been suggested to regulate TAZ phosphorylation and its nuclear localization (Zhang et al., 2016), as a scaffold for YAP-binding proteins. The Notch signaling pathway is simpler than the Hippo-YAP/TAZ pathway, as Notch/NICD acts as a “membrane-tethered” transcriptional co-activator, without the complex cascade of kinase-dependent steps as in YAP/TAZ regulation. Nonetheless, NICD is still regulated by phosphorylation events, for example to limit prolonged pathway activity, through the action of Mastermind and CycC:CDK8 (Fryer et al., 2004). It is thus possible that Baz/Par3 may regulate NICD phosphorylation and nuclear localization as well.

Generally, the strength of activation of target gene expression depends on NICD stability within the resulting activator complexes, which also contain histone modifying/remodeling proteins, and accessibility of enhancers of any given target gene (Borggrefe and Oswald, 2009; Bray, 2016). N-signaling acts in many different contexts and can activate many distinct target genes, and as such the presence of context-determining factors will determine whether an enhancer is primed to react to a specific signal, which may result in a gene having different responses towards the same signals (Borggrefe and Oswald, 2009). Our data suggest that Baz/Par3 acts rather generally on NICD mediated target gene activation, as we see effects on Notch target expression in multiple tissues and on different target genes. It is likely that Baz/Par3 is directly or indirectly involved in promoting NICD association with Su(H), and the formation of the Su(H)-NICD-Mam complex. The precise function of Baz/Par3 in the context of NICD regulation will require transcription specific assays to be explored in future studies.

## Materials and Methods

### Drosophila genetics and strains

*w-IR* (*w^JF01540^*, P{TRiP.JF01540}), *baz-IR-1* (*baz^JF01078^*, P{TRiP.JF01078}), *baz-IR-2* (*baz^JF01079^*, P{TRiP.JF01079}), *aPKC-IR* (*aPKC^JF01966^*, P{TRiP. JF01966}), *UAS-mCD8RFP, baz^4^ FRT9-2* and *baz^815-8^ FRT9-2* fly stocks were obtained from Bloomington Drosophila Stock Center. *baz^815-8^* was recombined with FRT19A and then used for *baz^815-8^* mutant clonal analysis with FRT19A MARCM stock (genotype: *w, tub-gal80, hs-flp, FRT19A; act-Gal4, UAS-mCD8RFP / CyO*). *UAS-numb-GFP* was a gift from Bingwei Lu (Song and Lu, 2012).

*N^55e11^/baz^4^* and *N^55e11^/baz^815-8^*were generated by intercrossing via several crosses: *ubi-baz-mCherry* (Bosveld et al., 2012) was recombined with *Sb^1^* to generate *ubi-baz-mcherry Sb^1^* flies. *baz^4^*(or *baz^815-8^*) males with *ubi-baz-mCherry Sb^1^*which rescued *baz^-^* mutations (*baz^4^* or *baz^815-8^*; *ubi-baz-mCherry Sb^1^*/+) were crossed to *N^55e11^*/*FM7* females to generate *N^55e11^/baz^4^* or *N^55e11^/baz^815-8^*females (selected against *Sb^1^* phenotype).

*UAS-NICD* (Zaffran and Frasch, 2000) flip-on clones were analyzed and induced together with either *UAS-GFP*, *baz-IR1*, or *baz-IR2*, *UAS-numbGFP*, and *UAS-Su(H)Ank* (full genotypes listed in the figure legends). Third instar larvae were dissected 44-49 hours after clone inductions (which were induced via brief heat-shocks). Control samples from the same set of experiments were heat-shocked, processed and stained together to reduce experimental variations.

### DNA constructs

The *baz*-pENTR was kindly provided by Tony Harris (McKinley et al., 2012). Gateway cloning (Invitrogen) recombined pPWF (from Drosophila Genomics Resource Center) for *UASP-baz* (C-terminal 3X Flag tagged *baz*). The construct was sequenced with *baz* oligos to confirm correct recombination insertion and the Flag tag sequence. Transgenic flies were injected with *UASP-baz-Flag* ppWF plasmid by BestGene.

A fusion of *D. melanogaster* Su(H) and the Notch Ankyrin repeats was designed along the same principle as X-Su(H)-ANK (Wettstein et al., 1997). *Su(H)* coding sequence was cloned from GH10914 (DGRC Stock 5973) into pAc5.1/V5-His A (Invitrogen) using the following primers: Fwd: ACCGGAATTCACCatgaagagctacagccaatt and Rev: ACGCGGATCCggataagccgctaccatgac to remove the stop codon. The sequence encoding Notch Ankyrin repeats were cloned from pAc5.1/N^IC^-HA (gift of Ram DasGupta) using the following primers:

Fwd: ACGCGGATCCgatgtggatgcacgtggacct and

Rev: CTTTGCGGCCGCtcacagatcttcttcagaaataagtttttgttcatgctcgtccagcagcctga to introduce a stop codon and myc epitope. The Su(H)-ANK-myc sequence was then subcloned into pUASt-AttB, (and injected into UAS-AttP flies (RRID:BDSC_9724) by BestGene. Multiple transgenic lines were sequenced and phenotypically tested to confirm insertion of the UAS-Su(H)-ANK-myc sequence.

### Immunohistochemistry and microscopy

The following antibodies were used in the study: guinea pig Anti-Baz 1:200 (Blankenship et al., 2007), rabbit anti-Dlg 1:3000-4000 (Izaddoost et al., 2002), mouse anti-Wg 1:10-20 (Brook and Cohen, 1996), mouse Anti-Cut 1:20-1:40 (Blochlinger et al., 1990), rabbit anti-RFP 1:300 (Rockland Immunochemicals), rabbit anti-Galactosidase 1:2500 (Cappel). Mouse anti-Cut and anti-Wg antibodies were obtained from Developmental Studies Hybridoma Bank at the University of Iowa.

Adult wings and nota were imaged with conventional Zeiss Axioplan microscope. Immuno-fluorescent samples were imaged with Leica TCS SP5 or SP8 confocal microscope. Images were projected and quantified with Fiji-Image J. Cut expression levels were measured in cell nuclei marked by Hoechst staining. *baz*^815-8^ mutant cells were outlined by RFP expression. Only first row of *baz*^815-8^ mutant cells right at D/V border were measured for “average Cut channel pixel intensity” minus “average background pixel intensity outside Cut expression domain” (similar to Figure S4D1) for each nucleus. Neighboring control A/P border cells were also measured for “average Cut pixel intensity” minus “average background pixel intensity”.

## Data visualization and Statistics

Data were plotted via Microsoft Excel, LibreOffice Calc. Fisher’s exact test (two tailed) and student t-test (unpaired) were used for statistical analyses to calculate probabilities of null hypothesis with the web version of Graphpad (Fisher’s exact test) and scipy.stats (t-test).

## Cell Culture and co-immunoprecipitation

NICD-HA pAc and myc-Hpo pAc were generous gifts from Ramanuj DasGupta and Duojia Pan (Wu et al., 2003). Baz-myc pAWM was generated from pAWM gateway vector (Drosophila Genomics Resource Center) and *baz* pEntr 2B (a gift from Tony Harris).

1mg of NICD-HA pAc and Baz-myc pAWM or myc-Hpo pAc plasmids each were transfected into *Drosophila* S2 cells in a 6-well plate with Qiagen Effectene. Cells were lysed one day after transfection with IP buffer (20 mM HEPES pH 7.5, 100 mM NaCl, 0.05% Triton X100, 1 mM EDTA, 5 mM DTT, 10mM NaVO4, 10% glycerol and protease inhibitor cocktail). The cell lysates (1mg of total protein) were pre-cleared by incubating with protein A-sepharose beads (Thermo Scientific 20333) for 1 hour at 4°C, followed by centrifugation to remove the beads. Mouse anti-Myc antibody (sc-40, Santa Cruz technology) was used at 1/1000 dilution for immunoprecipitation. The A-sepharose beads and antibody were incubated together to form pre-immuno-precipitate complexes at 4°C for 1 hour, then incubated with lysates. Myc-tagged protein and associated proteins were captured by protein A-sepharose/antibody complexes after overnight incubation at 4°C. Samples were washed 4 times with IP buffer. Immunoprecipitates were resuspended in SDS sample buffer, boiled for 5 min, separated by SDS-PAGE, and transferred to nitrocellulose membrane for immunoblotting. The blots were incubated with mouse anti-myc (Santa Cruz technology) or rat anti-HA (Roche) at 1:1000, subsequently with anti-mouse or anti-rat IgG-HRP. Proteins were then detected by enhanced chemiluminescence kit (Millipore).

## Acknowledgements

We are grateful to Kwang-Wook Choi, Kyung-ok Cho, Ramanuj DasGupta, Tony Harris, Duojia Pan, Bingwei Lu, and Jennifer Zallen for *Drosophila* strains and antibodies, and to Cathie Pfleger for helpful suggestions on the manuscript. We also thank the Bloomington Drosophila Stock Center for fly strains, and the Developmental Studies Hybridoma Bank at the University of Iowa for antibodies. We would also like to thank the ISMMS Microscopy CoRE, where confocal microscopy was performed and was in part supported by the Tisch Cancer Institute P30 CA196521 grant from the NCI. This work was supported by National Institutes of Health grants R01 EY013256 and R35 GM127103 (to MM).

