## Supplemental Data for "Par3/Bazooka binds NICD and promotes Notch signalling during *Drosophila* development"

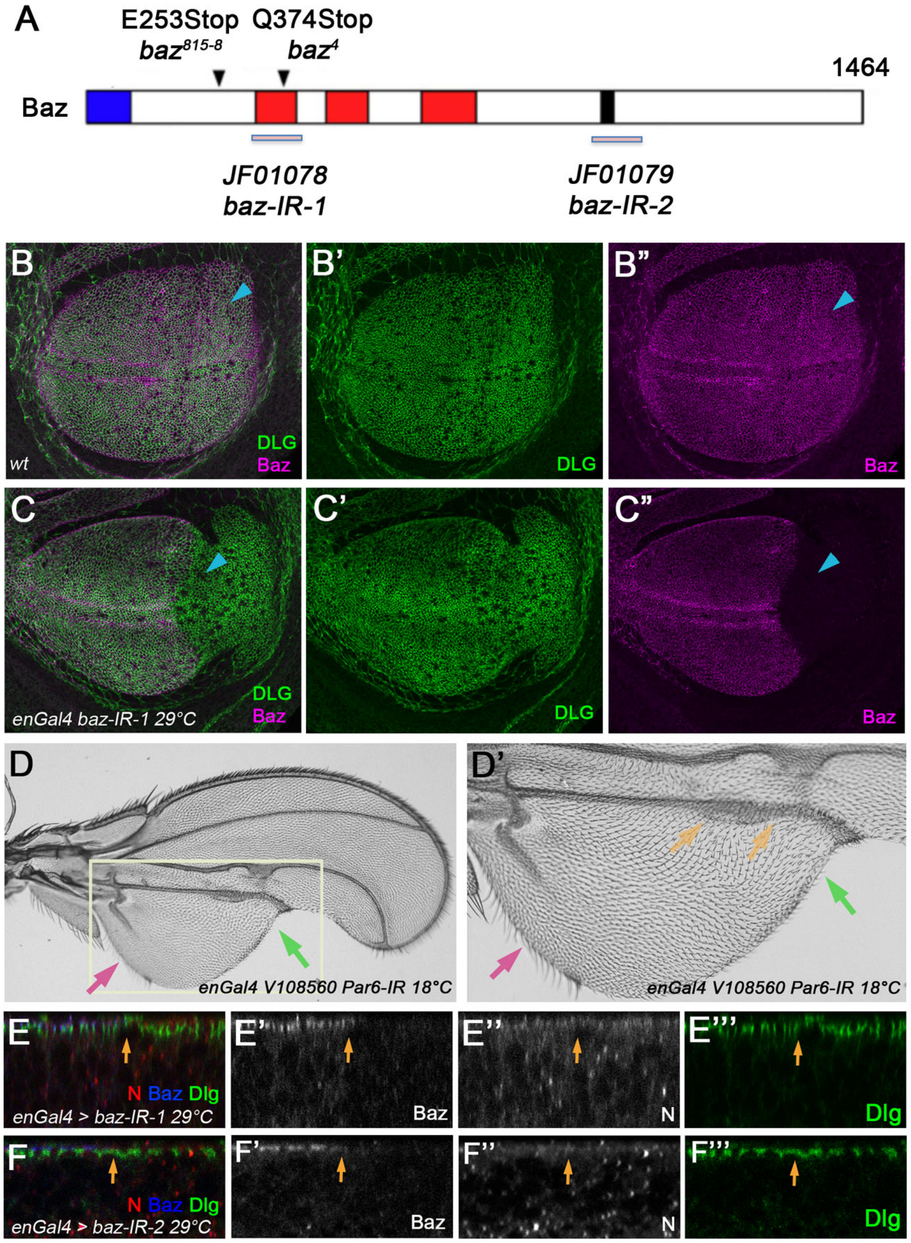


**Figure S1 (supplement to Figure 1):**

***baz reagent* characterization and the effect of *Par6* knockdown in the wing.**

(A) Graph illustrating the positions and characteristics of the alleles $baz^4$, $baz^{815\text{-}8}$, and RNAi transgenes *baz-IR-1* and *baz-IR-2*.

(B-C) Third instar larval wing discs stained for Dlg (green) and Baz (magenta). (B) Both Dlg and Baz are expressed in the anterior and posterior compartments in wildtype wing discs. (C) In the *baz-IR-1* knockdown condition (driven by *enGal4*), Dlg remained similarly expressed and localized in both anterior and posterior compartment, suggesting that overall tissue architecture and epithelial polarity is not affected. Baz was markedly reduced if not completely lost in the posterior compartment. Genotype: *UAS-dcr2; en-Gal4 / baz-IR-1 at 29°C*.

(D-D') The Gal4-UAS system is temperature sensitive with higher temperature leading to higher expression of UAS-transgenes. *Par6-IR* was expressed in the posterior compartment at lower temperature 18°C to reduce tissue loss and allow analysis. *Par6* knockdown caused loss of wing cells/tissue (green arrows). In regions where tissue loss was not predominant (magenta arrows), wing margin was relatively normal, suggesting N-signaling was not directly affected. Genotype: *en-Gal4/V108560* at 18°C.

(E-F) Notch levels and localization were not significantly altered in *baz-IR* expressing cells.

In the posterior compartment of *enGal4* driven *baz* knock-down wing discs, we did not detect differences in Notch protein levels and/or localization as compared to the anterior (the posterior compartment is marked by loss of Baz; blue in E and F, and monochrome in E' and F'; the A/P-compartment boundary is highlighted by orange arrow in these panels). Baz-staining was detected strongly near adherens junctions in the anterior, and was missing in posterior *baz-IR* expressing region (E, E' with *baz-IR-1*; or F, F' with *baz-IR-2*). Notch (red in E, F, monochrome in E", F", as detected with an anti-NICD antibody) staining and localization was not significantly altered in *baz-IR-1 or baz-IR-2* expressing regions. Dlg (green) expression and localization was indistinguishable between the two compartments, suggesting the epithelial architecture and A/B polarity remained unaffected. Genotypes: (E) UAS-dcr2; *en-Gal4 / baz-IR-1 at 29°C.*

(F) *en-Gal4 / baz-IR-2 at 29°C*.

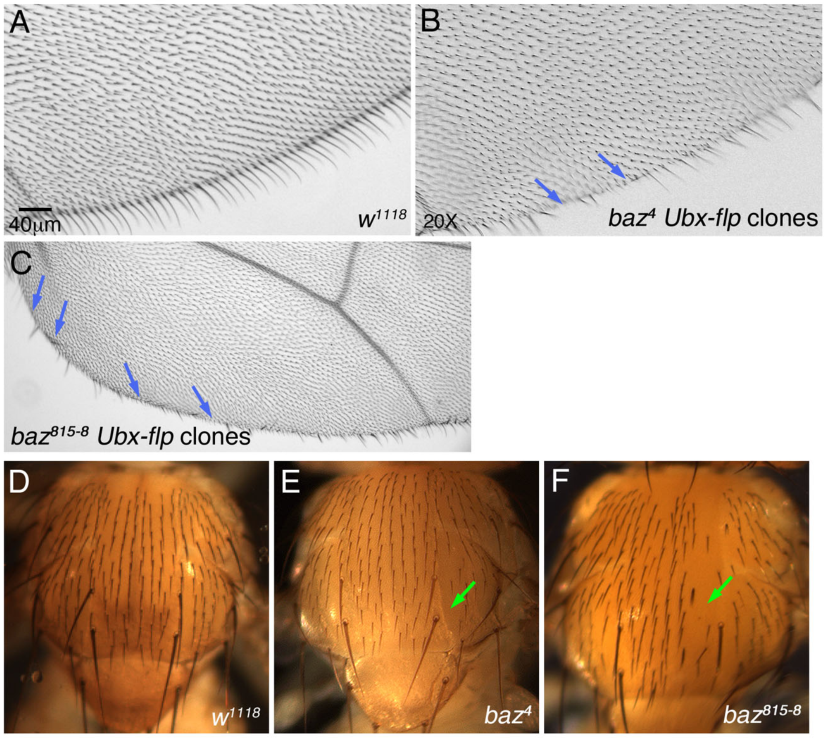


**Figure S2 (supplement to Figure 1):**

**Clonal *baz*⁻ mutant phenotypes in adult wings and nota.**

(A-C) Wings of wild-type (A) $baz^4$ (B) or $baz^{815-8}$ (C) mutant clones (unlabeled) induced with the Flp/FRT-system [56] via *Ubx-flp*. *Ubx-flp* tends to generate many small clones. Margin loss was observed in wings containing $baz^4$ (B) or $baz^{815-8}$ (C) mutant clones.

(D-F) Nota from wild-type (D) or containing $baz^4$ (E) or $baz^{815-8}$ (F) mutant clones (also induced with *Ubx*-driven Flipase), displaying sensory bristle loss and double bristle phenotypes (E, F), consistent with previous publications [21]. Genotypes: (B, E) $baz^4$ *FRT9-2 / ubi-GFP FRT9-2; Ubx-flp/+*. (C, F) $baz^{815-8}$ *FRT9-2 / ubi-GFP FRT9-2; Ubx-flp /+*.

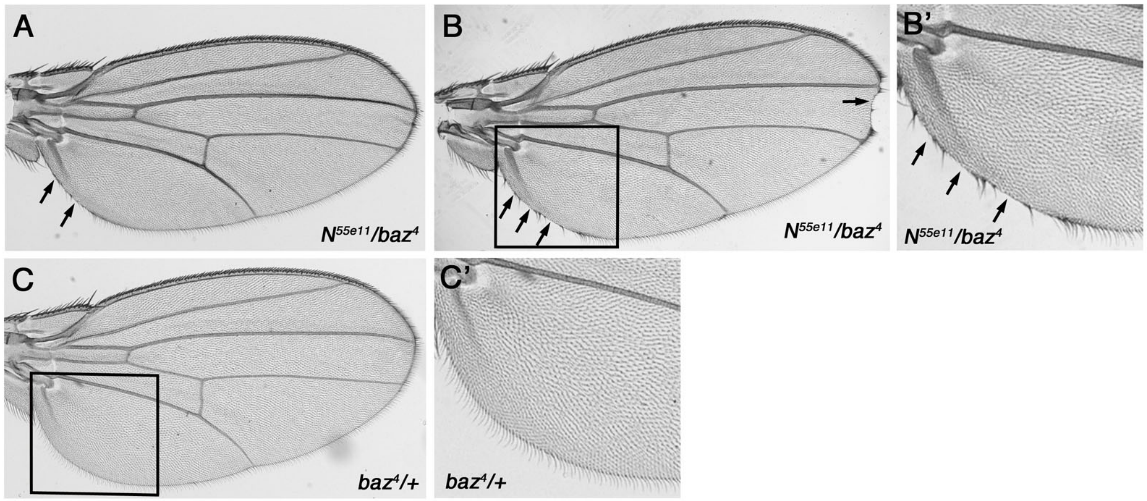


**Figure S3 (supplement to Figure 2):**

**Genetic interactions between *Notch* and *baz* alleles.**

(A-B) $N^{55e11}/baz^{4}$ double heterozygous wings displaying increased wing margin loss as compared to $N^{55e11}$/+ alone (B, arrow). Note also that in addition to the distal notching, the proximal-posterior margin area displayed always a margin loss even when distal margin had no notching defects (A); see also (B and B', arrows). (B') Higher magnification view of boxed area in B. Compare to main Figure 2 for different *baz* allele and quantifications in main Figure 2F-G.

(C) $baz^{4}$/+ single heterozygous wings showed no phenotypes and a fully wild-type wing margin. (C') Higher magnification of boxed area in C. Quantifications in main Figure 2F-G.

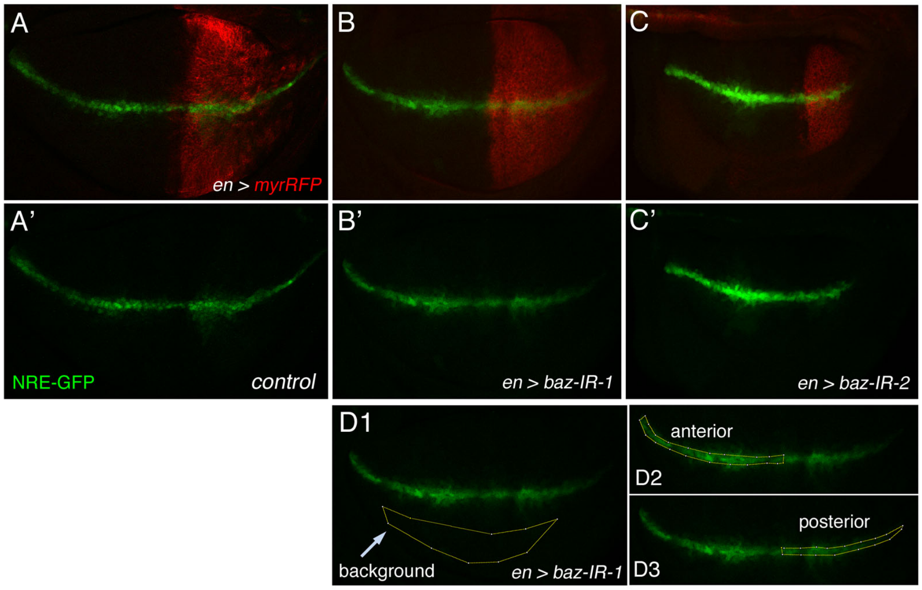


D4

$$\text{intensity ratio} = \frac{(\text{average pixel intensity of the posterior - average pixel intensity of the background})}{(\text{average pixel intensity of the anterior - average pixel intensity of the background})}$$

E

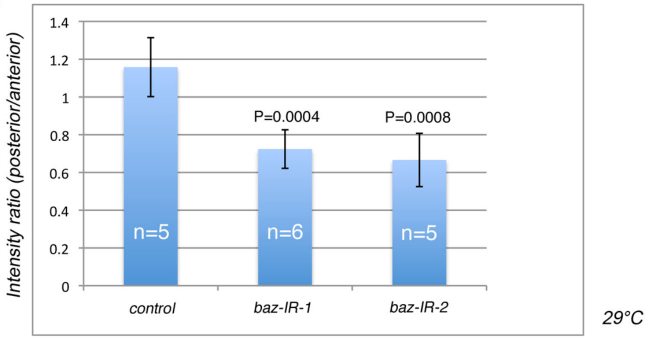


**Figure S4 (supplement to Figure 3):**

***baz-IR* knock-down reduced expression of Notch response element (NRE) reporter gene.**

(A-C) *NRE-GFP* is expressed in a Notch-signaling dependent fashion at the dorsal/ventral boundary in wild-type wing discs [31], a pattern mimicking N-signaling activation and endogenous Notch target gene expression, e.g. *wg* and *cut* expression patterns. Myr-RFP alone (A, control), myrRFP and *baz-IR*-1 (B) myrRFP and *baz-IR*-2 (C) were expressed in the posterior compartment under *en-Gal4* driver control (MyrRFP marking cells expressing *en* in posterior compartment). Note that *NRE-GFP* expression was reduced in the posterior compartment (see quantification in E).

(D1-4) The panel illustrates how anterior and posterior expression levels were quantified. D1: average background signal level was measured, inside yellow outlined region not showing any specific expression. D2: average anterior *NRE-GFP* levels, measured in outlined region. D3: average posterior *NRE-GFP* levels, measured in outlined region. Anterior / posterior border was defined by MyrRFP expression. D4: formula for posterior/anterior intensity ratio calculation with deducted of background signal.

(E) Quantification plot of *NRE-GFP* expression levels as posterior/anterior intensity ratio in the different genetic backgrounds. The posterior/anterior intensity ratio decreased significantly under *baz-IR* conditions with $p = 0.0004$, or $0.0008$ respectively for *baz-IR-1* or *baz-IR-2*. Student t-test was used for the $p$ values.

Genotypes: (A) *UAS-dcr2; en-Gal4, UAS-MyrRFP, NRE-EGFP/CyO at 29°C.*

(B, D, E): *UAS-dcr2; en-Gal4, UAS-MyrRFP, NRE-EGFP/+; baz-IR-1/+ at 29°C*.

(C, E): *UAS-dcr2; en-Gal4, UAS-MyrRFP, NRE-EGFP/+; baz-IR-2/+ at 29°C*.

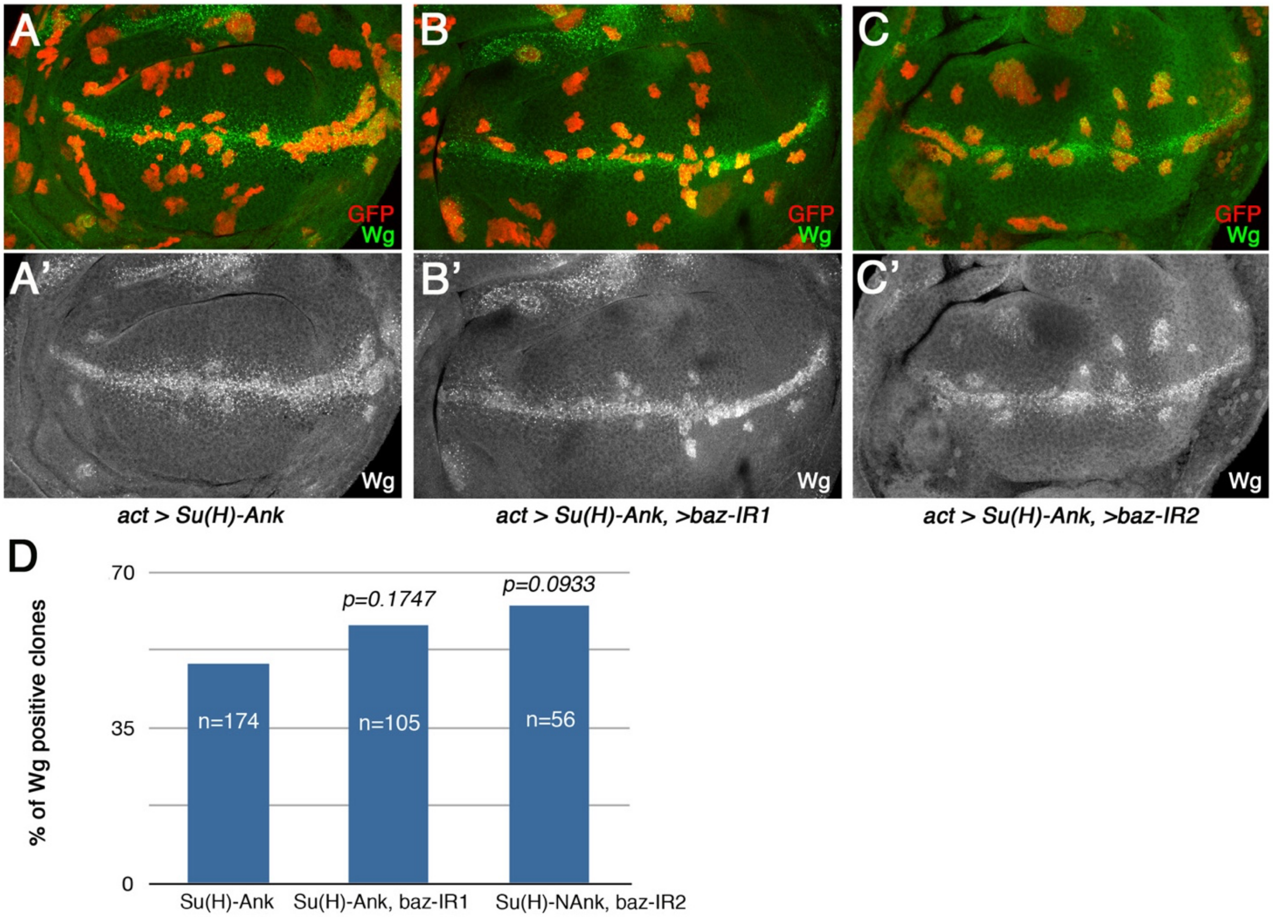


**Figure S5 (supplement to Figure 5)**

**Knockdown of *baz* does not affect Su(H)-Ank mediated Notch target gene induction.**

Su(H) and Notch Ank repeat fusion protein (Su(H)-Ank) was previously shown to activate Notch signaling (Wettstein et al., 1997 and Collu et al. 2012). We tested whether reduction of *baz* levels (via RNAi) could interfere with the induction of the target gene *wg*.

(A) Control confocal projection of wing disc showing clonal induction of cells expressing Su(H)-Ank (using the *act-Gal4* flip-on technique) with GFP as clonal marker. Wg expression is normally restricted to the dorsal/ventral boundary (detected with anti-Wg antibody). Note that Su(H)-Ank clones can express the target gene *wg* within the wing pouch area, generally near endogenous wg expression domain. (A') showing the Wg monochrome. Genotype: *hs-Flp, UAS-dcr2/Y; act5c>y>Gal4, UAS-GFP/UAS-Su(H)-Ank.*

(B-C) Confocal images of clonal expression of Su(H)-Ank together with *baz-IR-1* (B) or *baz-IR-2* (C)(see Suppl. Figure 1 for location of the independent *baz* IRs within the *baz* gene). Again, note that Su(H)-Ank clones can express the target gene *wg* within the wing pouch area. (B'-C') showing wg expression in monochrome. Genotype in B: *hsflp, UAS-dcr2/Y; act5c>y>Gal4, UAS-GFP/UAS-*

*Su(H)-Ank*; *baz-IR-1/+*. Genotype in C: *hs-Flp, UAS-dcr2/Y; act5c>y>Gal4, UAS-GFP/UAS-Su(H)-Ank*; *baz-IR-2/+*.

(D) Quantification of repeat experiments of genotypes shown in (A-C). Note that there was no reduction in clones inducing Wg expression in wing discs co-expressing *baz-IR-1* or *baz-IR-2* with Su(H)-Ank, as compared to Su(H)-Ank alone. This suggested that *baz* is not required for Su(H)-Ank mediated transcriptional activation of the Notch target *wg*. Differences between Su(H)-Ank and Su(H)-Ank/*baz-IRs* were not significant (Fisher's exact test).

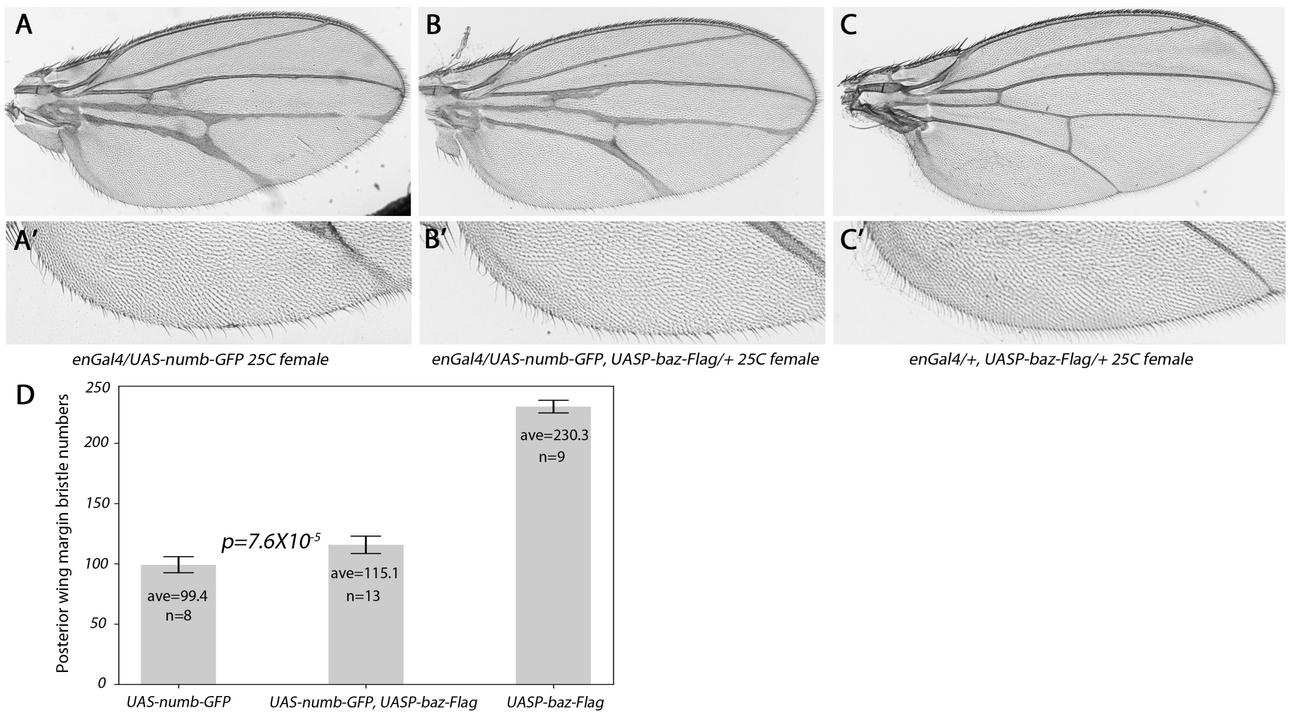


**Figure S6 (supplement to Figure 7):**

***numb* and *baz* genetically interact during Notch signaling mediated wing patterning.**

(A-C) *UAS-numb-GFP* (GFP tagged version of Numb) was expressed in the posterior compartment of the wing via *enGal4* driver control with or without *UASP-baz-Flag* (at 25°C). Note that here and in all other wing experiment only female flies were used to avoid possible variation between males and females. (A, A') Expression of *numb-GFP* caused wing margin loss and thickening of veins in the posterior compartment (both features indicate an inhibition of Notch signaling). Genotype: *enGal4/UAS-numb-GFP*. (B, B') Expression of *numb-GFP* together with *baz-Flag* caused a partial rescue of wing margin cell loss and vein thickening. Genotype: *enGal4/UAS-numb-GFP; UASP-baz-Flag/+.* (C, C') Control wing samples expressing *UASP-baz-Flag* only. Such wings had a fully wild-type appearance with approx. 230 margin bristle cells. Genotype: *enGal4/+; UASp-baz-Flag/+*. (D) Quantification illustrating the effects on posterior wing margin bristle numbers in the genotypes shown in (A-C). Note that co-expression of Baz significantly reduced wing margin bristle loss caused by Numb overexpression. The number of margin bristles in UASp-Baz-Flag is equivalent to wild-type.
